# Latitudinal Patterns in Immunogenetic and Microbiome Diversity in two anuran species: *Bufo bufo* and *Bufo spinosus*

**DOI:** 10.64898/2026.07.10.737765

**Authors:** Eleonore Susi, Zhenzhen He, Filip Thörn, Patrik Rödin-Mörch, Niki Chondrelli, Barbora Thumsova, Jaime Bosch, Anssi Laurila, Jacob Höglund, Maria Cortazar-Chinarro

## Abstract

Evolutionary and demographic processes such as selection, drift and migration shape the genetic variation of populations. Genetic diversity is often lower in populations toward higher latitudes. This decrease potentially threatens their survival as several factors are putting more pressure on the populations, including the spread of infectious diseases. In this study, we combined whole-genome re-sequencing with MHC class II genotyping and skin microbiome profiling in *Bufo bufo* and *B. spinosus,* two closely related European toad species. We investigated the underlying immunogenetic and microbial variation resulting from different demographic histories and environmental conditions to identify their potential impact on infection outcomes in these two species. We found lower immunogenetic diversity in *B. bufo* compared to *B. spinosus*, with highly significant differences in genes related to adaptive and innate immunity. We found lower overall MHC class II diversity and skin microbiome diversity at the species level in *B. bufo*, compared with *B. spinosus*. In contrast, at the individual level, *B. bufo* showed higher MHC allelic diversity and greater diversity in the core skin microbiota than *B. spinosus*. Together, our findings suggest that divergence in immunogenetic background and host-associated microbial communities may underlie differences in susceptibility to emerging infectious diseases. This integrative framework provides new insight into how host genetics and microbial communities jointly influence disease outcomes across environmental gradients.

## Introduction

Historical events such as colonization processes can profoundly influence patterns of genetic variation [1]. Repeated founder events—where new populations are established from a few individuals—result in substantial losses of genetic diversity [2–4]. During the Pleistocene, much of Northern Europe was glaciated, forcing most extant species to retreat to southern refugia, and subsequent post-glacial colonization during the Holocene eroded genetic variation through successive founder effects [5,6]. Consequently, populations at high latitudes in Europe typically exhibit reduced genetic diversity. At a broader scale, genetic diversity declines from tropical to polar regions across a broad range of taxa, including plants and animals [7,8]. The decline in genetic variation is crucial, as genetic diversity underpins a population’s capacity to respond to environmental change and influences evolutionary potential, local adaptation, and long-term population persistence [9–11]. In amphibians, both overall genetic variation and immunogenetic diversity decrease with increasing latitude in European populations [11–14]. Currently, amphibians face population declines driven by global change and anthropogenic pressures, including habitat fragmentation and emerging infectious diseases such as the chytridiomycosis caused by the fungus *Batrachochytrium dendrobatidis (Bd*) and ranavirosis caused by *Ranavirus* (*Rv*) [15,16]. Population-level genetic variation in the immune system can be of considerable importance for population survival [17–19].

The major histocompatibility complex (MHC) is a multi-gene family involved in the recognition and presentation of pathogen-derived antigens to immune cells, thereby initiating adaptive immune responses [20]. MHC class II molecules present peptides derived from extracellular pathogens and play a central role in shaping CD4 T-cell–mediated immunity [20]. Diversity at MHC class II loci has been associated with variation in susceptibility to both fungal and viral infections across multiple vertebrate systems, including fishes, amphibians and mammals [21–23]. In amphibians, specific MHC class II alleles and supertypes have been linked to resistance against the fungal pathogen *Bd* [17,24,25] and viral pathogens [19]**Error! Bookmark not defined.**. Similarly, Fu [18] found that chytrid-resistant amphibian species possessed distinct MHC class II variants compared with susceptible species, with evidence of positive selection on peptide-binding residuals, suggesting an important role of MHC class II molecules in defending against *Bd* infection. Thus, reduced immunogenetic diversity at higher latitudes may impair their ability to combat hypothetically northward-spreading epidemics like *Bd* and *Rv* infections [26–29].

All multicellular organisms host commensal microbiomes that protect against pathogen invasion through direct mechanisms, such as antimicrobial compound production, and indirect ones, including host immune response modulation [30–32]. The skin and its associated microbiota constitute the first line of defence against environmental pathogens [34,35], with commensal bacteria suppressing invaders through metabolic interference and facilitating host adaptation to specific ecological challenges [33,34]. In amphibians, this protective role is particularly critical, as the skin represents the primary entry point for pathogens such as *Rv* and *Bd* [35,36]. The amphibian skin microbiome thus functions as a key component of the extended immune system, complementing host genetics and innate immune defences [37], and its composition and diversity have been extensively studied in relation to pathogen resistance [37–40]. Nevertheless, characterizing microbial communities across species and populations in relation to immune genetic diversity remains a critical gap in understanding how microbiome variation relates to infectious disease dynamics in wild animals.

The common toad *Bufo bufo (Bb)* and the spiny toad *B. spinosus (Bs)* are two closely related European bufonids [41], where emerging evidence suggests interspecific variation in their susceptibility or physiological responses to infectious diseases [42,43]. *Bs* shows high susceptibility and mortality to both *Bd* and *Rv* in the Iberian Peninsula [44,45], whereas die-offs of *Bb* have not been reported. Experimental studies have demonstrated that *Bd* induces mortality in *Bb* [46], particularly in individuals from northern populations with lower immune genetic diversity [47]. It is important to consider that individuals in laboratory infection studies may exhibit reduced microbiome diversity relative to wild populations, a factor that could potentially contribute to the higher mortality rates observed under experimental conditions [48]. Despite these experimental findings, mass mortality events have not been reported in wild *Bb* populations in Sweden, where *Bd* infection prevalence remains low (∼3.4%; [49]) compared to substantially higher levels observed for *Bs* (∼33%; [42]). This may reflect that *Bd*-associated mortality predominantly occurs at the end of metamorphosis, which could lead to the underdetection of mortality events in natural populations [50]. While genetic and immunogenetic mechanisms of disease susceptibility have been investigated primarily in *Bb*, studies on *Bs* have largely focused on population declines and disease impacts, with less attention to the underlying mechanisms that may explain its high susceptibility to emerging pathogens [17,51,52]. Comparative studies of closely related amphibian species have revealed contrasting disease outcomes but have focused mainly on transcriptomic responses rather than immunogenetic variation at MHC loci [53]. Similarly, although host-associated microbiomes are increasingly recognized as key components of defence, their role in species-specific susceptibility or resilience to pathogens remains poorly resolved [38,39]. These mechanisms are especially pertinent given the anticipated northward expansion of infectious diseases, including *Bd* and *Rv*, as a consequence of global climate change [54], particularly where amphibian populations exhibit lower genetic diversity [14]. We generally expect that interspecific differences in immune responses contribute to the lower observed susceptibility in *Bb*, despite both species occurring under climatic conditions broadly suitable for *Bd* growth (i.e., higher latitudes in *Bb* and higher altitudes in *Bs*).

In this study, we used whole-genome re-sequencing to investigate and compare immunogenetic variation between *Bb* and *Bs*. We also characterized MHC class II exon 2 to identify the major diversity components that might contribute to susceptibility to infectious diseases. We analysed 16S rRNA bacterial profiles to define the core skin microbiome as bacterial taxa consistently detected across individuals, irrespective of species identity, which are likely to reflect either functionally important host associated microbes or persistently enriched taxa shaped by shared environmental exposure [55,56]. Focusing on the core microbiome provides insight into the most stable component of the skin microbial community and is therefore more likely to capture long term host–microbe associations than transient taxa that contribute only to overall diversity [56]. We hypothesize that *Bb* will exhibit lower immunogenetic diversity than *Bs*, consistent with the expected latitudinal gradient in genetic diversity, but show higher diversity in disease-related genes, such as the MHC class II, which could explain the different mortality in association to infectious diseases. Alternatively, this reduced immunogenetic diversity in *Bb* may be partially offset by a distinct skin microbiome composition—either through elevated microbial diversity or functionally relevant shifts in core bacterial taxa—contributing to enhanced overall resilience to infection. Integrating immunogenetic and microbiome variation between *Bb* and *Bs* provides a mechanistic framework for understanding their differential vulnerability to infectious diseases and offers new insights into the processes that may underlie the higher field mortality observed in *Bs*, despite its broader species level genetic variation.

## Methods

### Sample collection

Tissue samples and skin microbiome swabs were collected from breeding adults of *Bs* in Spain and *Bb* in Sweden. In Spain, toe webbing clips were obtained from 84 individuals across six populations in four national parks: Barranco Lugar (N = 10, Ordesa), Lago Ercina (N = 20, Picos de Europa), Laguna de los Pajaros (N = 12, Sierra de Guadarrama), Laguna Grande (N = 12, Sierra de Guadarrama), Majada de la Vacas (N = 15, Sierra Nevada), and Soportujar (N = 15, Sierra Nevada). In Sweden, samples were collected from three populations in Kalmar province: Stjärnamo (N = 15), Kindbäcksmåla (N = 15), and Påryd (N = 13; total N = 43; Figure S1; Table S1). Tissue clips were stored in absolute ethanol (100% EtOH) at −20 °C. Skin microbiome swabs (side, belly, back, and legs) were collected using sterile cotton swabs after sequential rinsing of each individual in sterile water (Millipore Milli-Q™; 5 min, then 2 min in a fresh container) to remove transient microorganisms, with fresh nitrile gloves used per individual to prevent cross-contamination. Swabs were stored on ice and subsequently preserved at −80 °C. At each site, 2 L water samples were collected as environmental controls, filtered sequentially through 0.7 µm and 0.2 µm membranes to remove large particles and capture microbial cells, respectively, alongside sterile water blanks, and stored at −80 °C until analysis.

### DNA extraction and library preparation

Genomic DNA was extracted from 30 individuals (*Bb*, n = 15; *Bs*, n = 15) using the DNeasy Blood and Tissue Kit (Qiagen), and libraries prepared with the Illumina DNA Prep Kit and sequenced on a two lanes on NovaSeq XPlus platform (150 bp paired-end; NGI SNP&SEQ Technology Platform, SciLifeLab, Uppsala, Sweden).

MHC class II exon 2 was characterized from 118 of 127 individuals (*Bb*: n = 43; *Bs*: n = 84; remainder excluded due to extraction/amplification failure) using six forward and eight reverse primers (Table S2) following Zeisset & Beebee [57]. Libraries were prepared as in Cortázar-Chinarro et al. [11], with gel-based size selection, purification (MinElute Gel Extraction Kit, Qiagen), and Illumina adaptor ligation (ThruPLEX DNA-Seq Kit, Takara Bio; Tables S3–S4; Additional Information 1), and sequenced on an Illumina MiSeq (PE300; NGI SNP&SEQ, SciLifeLab).

For 16S rRNA metabarcoding, DNA was extracted from 120 skin swabs (*Bb*: n = 43; *Bs*: n = 77; AllPrep DNA/RNA Kit, Qiagen), and the V4 region amplified following a two-step PCR protocol [37] (Tables S5–S6; Additional Information 2). Libraries were quantified (Quant-iT PicoGreen, Invitrogen), equimolarly pooled, and sequenced on an Illumina MiSeq (NGI SNP&SEQ, SciLifeLab).

### Data Processing

For the whole-genome re-sequencing, the adapter sequences were removed from the reads with Trimmomatic (v0.39) [58]. The reads were aligned to the common toad reference genome (GCF_905171765.1; NCBI RefSeq assembly) using the Burrows-Wheeler Alignment MEM (BWA; v0.7.18) [59]. Duplicates were marked with Picard (v3.3.0) MarkDuplicates [60]. SNP and indel calling were done using GATK (v4.5.0.0) [61], and genotyping was done using GATK GenotypeGVCFs. SNPs and indels were separated using GATK SelectVariants and further filtered with GATK VariantFiltration and Vcftools (v0.1.16) to remove any low-quality reads [62]. Description of filtering can be found in Table S7. We used the *Bb* reference genome annotations to manually extract all immune-related genes. We used Bedtools (v2.31.0) intersect to extract immune-related SNPs from the SNPs vcf file [63].

Raw 16S amplicon data were processed using the DADA2 R package [64]. (Additional Information 3). MHC sequencing data were processed by merging paired-end reads with FLASH v1.2 [65], followed by chimera removal, deduplication, pseudogene filtering, and individual sample assignment using AMPLISAS with default parameters [66] (Additional Information 4).

### Genome-wide immune gene analyses

We used pcadapt package [67] in R to identify the immune genes that are under putative selection in each species, following the protocol with default parameters on https://bcm-uga.github.io/pcadapt/articles/pcadapt.html to extract the genes under putative selection. Assuming that the PCA analyses with pcadapt indicated that PC1 explains the species differences, we identified the immune SNPs that showed the highest differentiation in PC1 and labelled them as outliers, representing genes under stronger putative selection. All other immune SNPs were treated as non-outliers, consistent with the assumption that they are evolving under weaker or more relaxed selection. We used Plink (v 1.90b.3n) [68] for PCA plotting, which was visualized in R (v4.4.3, The R Foundation for Statistical Computing, 2025) using the QQman package.

Nucleotide diversity of immune SNPs was calculated for both species using VCFtools (v0.1.16) [62] and SNPs were mapped to genes based on genomic coordinates using Bedtools intersect and Bb annotations (GCF_905171765.1; NCBI RefSeq). Per-gene nucleotide diversity was estimated as the mean π across constituent SNPs using tidyverse [69], and genes were assigned to eight immune-related Gene Ontology categories [70,71] (See additional Information 5).

### MHC II exon 2 and skin microbiome data analyses

#### MHC class II exon 2 diversity

Nucleotide diversity (π) and Watterson’s theta (θ_W_) were computed using DNAsp 6.0 [72] at three hierarchical levels: (1) between species, (2) between populations, and (3) between individuals (See Tables S8-10) to capture intra-individual variation between populations. We first counted the number of alleles for each individual. Then, we calculated π and θ_W_ based on all alleles present within an individual (Table S10). Due to the non-normal distribution of the data, which could not be resolved by transformation, we used a Wilcoxon rank-sum test to assess the effect of species identity on individual-level π and θ_W_ per site. We estimated MHC class II exon 2 allele frequencies based on both DNA and amino acid sequences identified in both species and subsequently visualized as pie charts generated with ggplot2 in R [73].

Phylogenetic trees of all MHC class II exon 2 alleles were constructed in MEGA 12 [74]. using maximum likelihood with 1000 bootstrap replicates. Substitution models were selected by BIC: JC+G and T92+G for *Bb* and *Bs* nucleotide alignments, respectively, and LG+G for amino acid alignments in both species.

#### Microbial diversity and core microbiome analysis

To verify that our sampling design minimized environmental cross-contamination, we compared microbiome composition between non-rarefied swab and environmental samples by PERMANOVA and PERMDISP tests using vegan package [75]. The environmental samples were removed before downstream analysis.

To evaluate bacterial community structure, we performed diversity analyses on both the full dataset and the core microbiome, defined as amplicon sequence variant (ASVs) with a relative abundance > 0.01% present in at least 70% of samples (microbiome package; [76]). Prior to downstream analysis, environmental cross-contamination was assessed via PERMANOVA and PERMDISP using the vegan package [75] after which, environmental samples were removed.

For full dataset diversity estimation, data were rarefied to 90% of the minimum sequencing depth using phyloseq [77]. Core microbiome diversity and composition were analyzed using the non-rarefied core-only dataset. Alpha diversity (Observed richness, Chao1, Shannon, and Simpson indices) was compared between *Bb* and *Bs* using Wilcoxon rank sum tests with Benjamini-Hochberg correction (rstatix package; [78]). Beta diversity was quantified using Jaccard, Bray-Curtis, and unweighted Unifrac distances. Community structure differences were tested using PERMANOVA (adonis2) and PERMDISP (vegan), and visualized via PCoA, NMDS, and circular hierarchical clustering using phyoseq [77], ape [79], and ggtree package [80].

#### Differential abundance and functional annotation

We performed differential abundance analyses on non-rarefied data (both full and core-only) using a consensus approach. ASVs were considered significantly differentially abundant only if they showed concordant effect directions (same log fold-change sign) and adjusted p-values < 0.05 in both LinDA [81] and DESeq2 [82]. Results were visualized using volcano plots (ggplot2; [73]) and heatmaps (ComplexHeatmap; [83]).

Finally, to identify putatively beneficial bacteria, we cross-referenced the genera of all differentially abundant and core ASVs against the Antifungal Isolates Database [84]. ASVs belonging to genera with documented inhibitory activity against *Batrachochytrium dendrobatidis* (*Bd*) were identified as putatively *Bd*-inhibitory taxa.

## Results

### Raw data processing

Raw data processing is summarized for the whole genome re-sequencing, MHC class II and 16s metabarcoding in Additional information 6.

### Genome-wide population structure

PCA revealed distinct population clustering of *Bs* for both genome-wide and immune SNP datasets (PC1 = 80.1% and PC1 = 78.5%, respectively; Figure 1). The populations follow a latitudinal geographic gradient, with the northernmost population (Picos de Europa) and the southernmost population (Sierra Nevada) exhibiting the greatest differentiation. In *Bb*, individuals clustered closely based on genome-wide SNPs, reflecting little population structure. Immune SNPs, however, displayed a more scattered pattern, suggesting differentiation at the individual level, though no distinct population clusters were evident.

**Figure 1.**
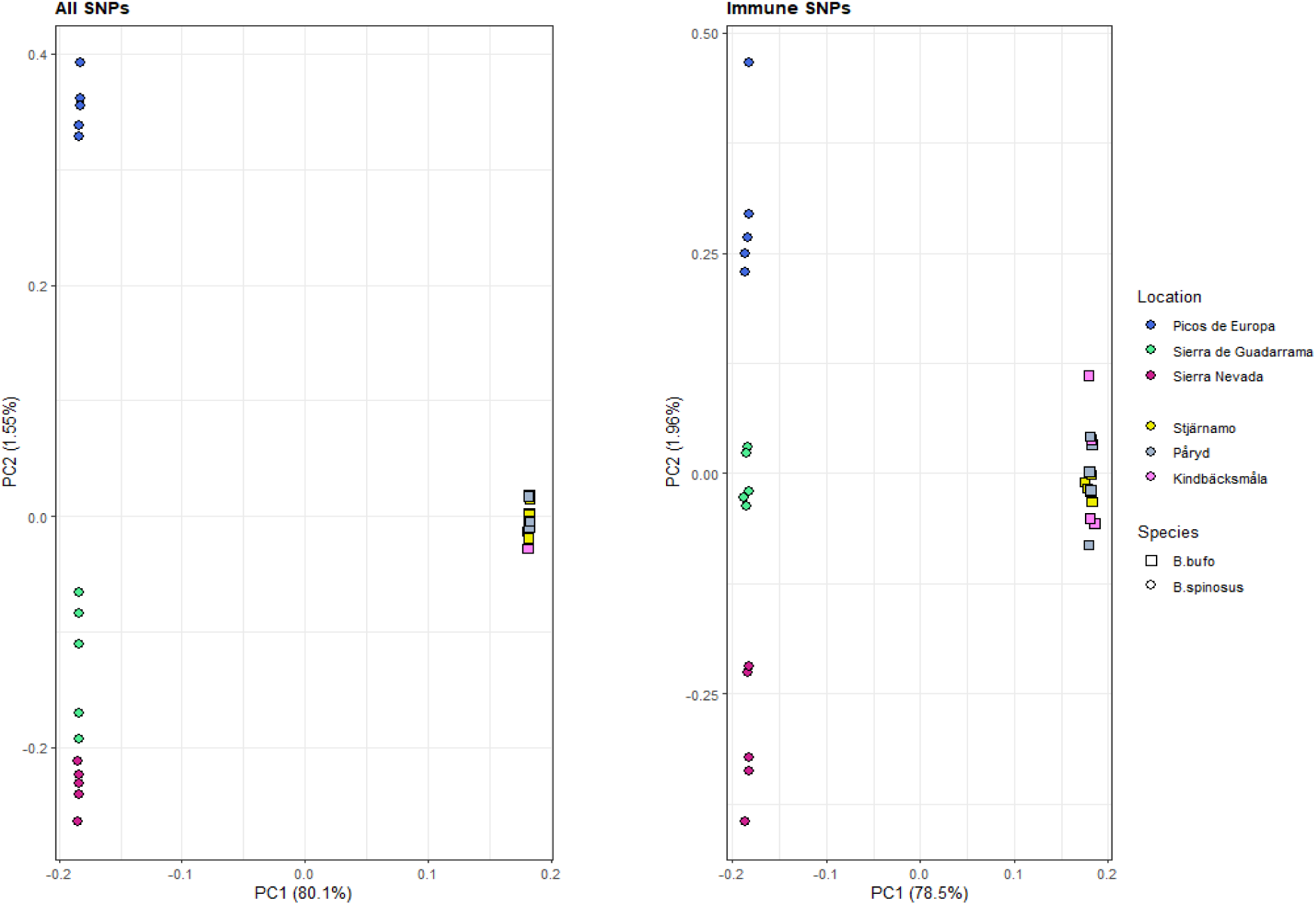
PCA of *B. bufo and B. spinosus* with their populations using genome-wide (All) SNPs and immune SNPs. Locations are indicated by colors with populations sorted from Northernmost to Southernmost. Species are indicated by shape.

### Genome-wide immunogenetic diversity

The pcadapt analysis identified 20,631 immune-related SNPs as outliers (*Bb*, N=15,614; *Bs*, N=10,964) and 67,988 SNPs as non-outliers (*Bb*, N=51,961; *Bs*, N=40,689). Among the immune-related SNPs outliers, 5,947 SNPs were shared between the two species, while 24,662 SNPs were shared among the non-outliers. Our results show highly significant differences between the outlier and non-outlier genes in *Bs* and *Bb* (Figure 2), with both species showing significantly lower nucleotide diversity in outlier genes. We also identified highly significant differences between *Bs* and *Bb* outlier genes and between *Bs* and *Bb* non-outlier genes. *Bb* showed lower nucleotide diversity in both outlier (*Bs*, mean π =0.057; *Bb*, mean π =0.026) and non-outlier genes (*Bs*, mean π =0.243; *Bb*, mean π =0.12), indicating an overall lower immunogenetic diversity in *Bb*.

**Figure 2.**
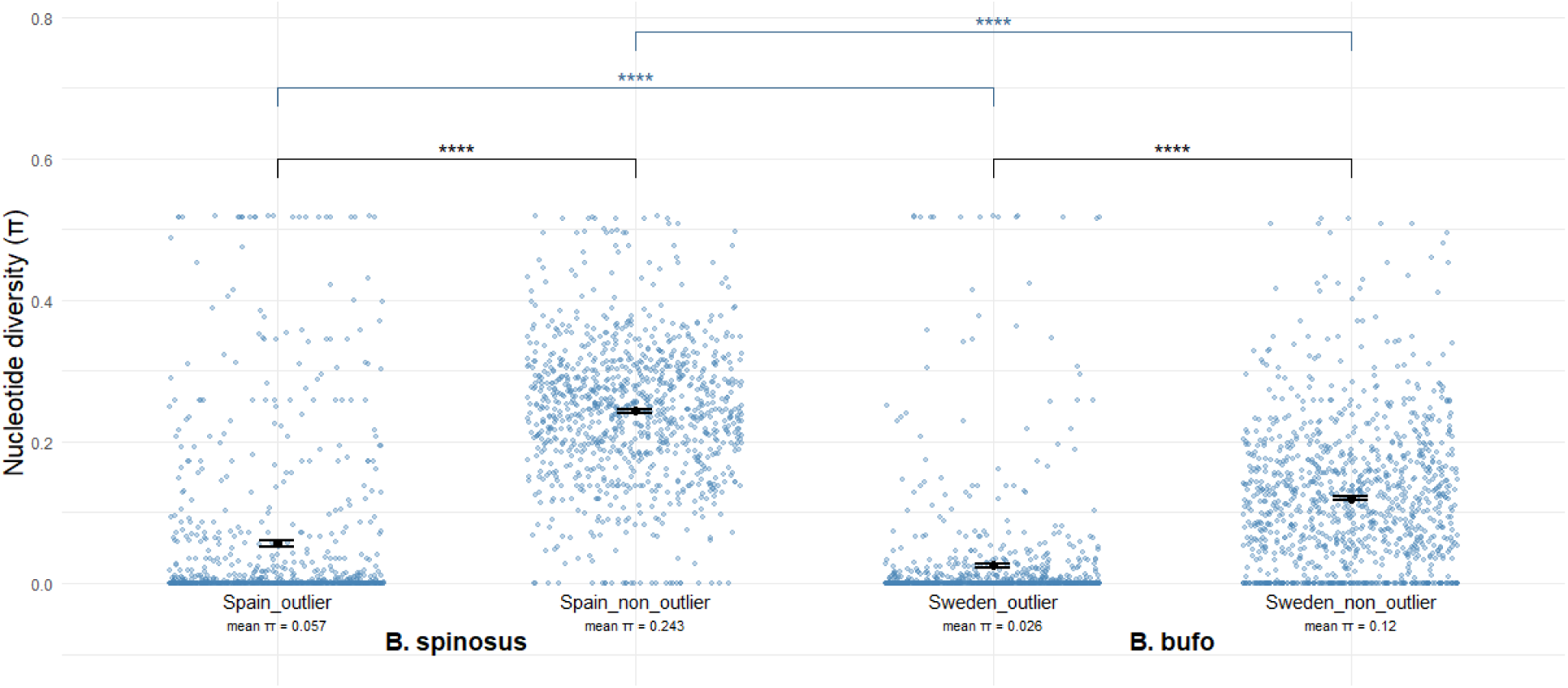
Nucleotide diversity of immune genes in *B. spinosus* and *B. bufo*. Significance was calculated between the outlier and non-outlier genes within species (black) and between the species (blue).

To identify more specific immunogenetic differences, we matched the immune SNPs to immune genes and grouped them into immune categories based on their main function. In total, 163 immune genes (*Bs*, N=155; *Bb*, N=161) were matched to immune-related categories (Figure 3). We found that *Bs* showed a higher mean π than *Bb* in all categories, with “Adaptive Immunity” and “Innate Immunity” both showing highly significant (p<0.001) differences between the species, and “Lymphoid and Organ Development”, “Cytokine and Interferon Response” and “Other” showing significant differences (p<0.005-0.05). Based on nucleotide diversity differences, *Bs* showed higher diversity in 3 characterized genes (C1S, LIF, TNFSF15) with differences ≥0.4, in 5 characterized genes (TRIM59, C8B, CD40, CD79B and LTB) with differences ≥0.3 and in 2 characterized genes (CD3E, and TNFRSF11A) with differences >0.2. Additional information can be found in the supplementary (Tables S11-14).

**Figure 3.**
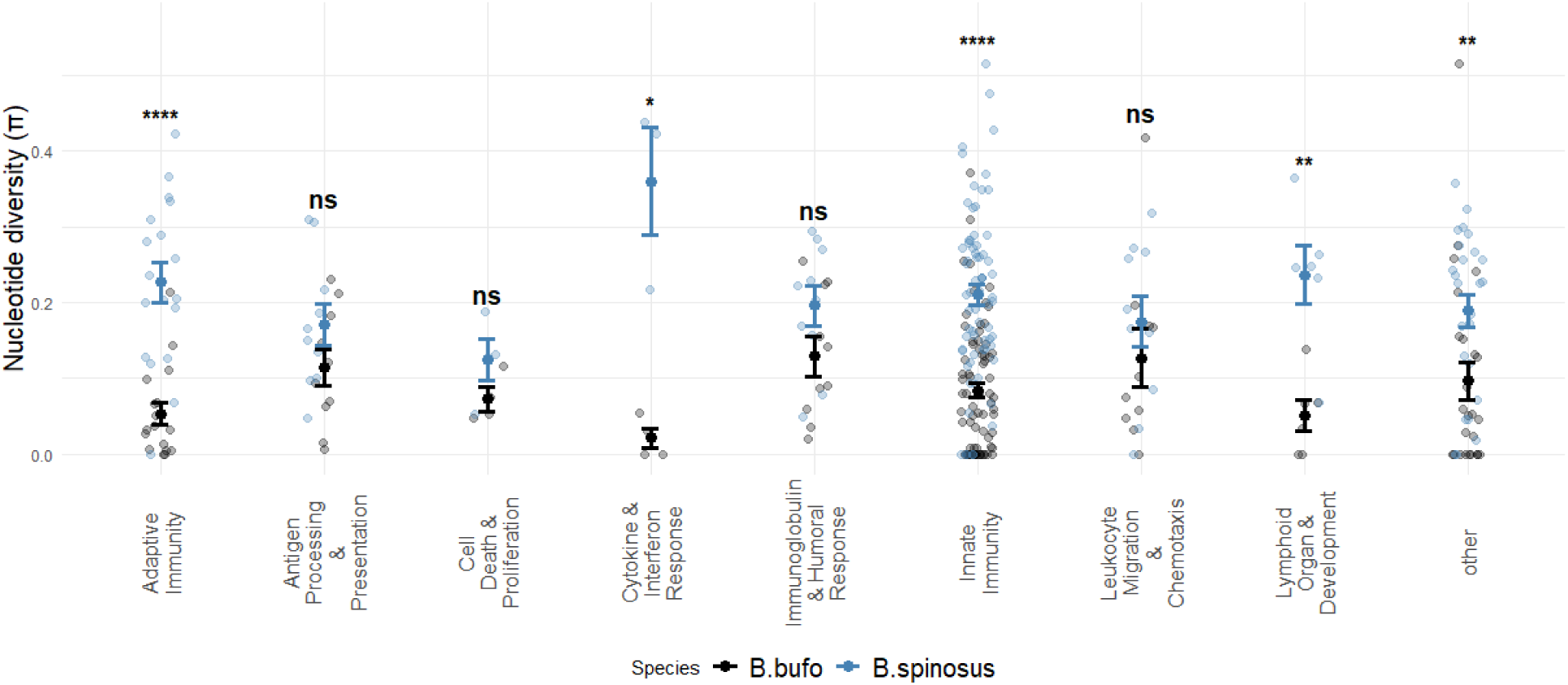
Nucleotide diversity of the immune genes based on immune categories in *B. bufo* and *B. spinosus*. The significance was calculated between the species.

### MHC class II exon 2 characterization

We identified 23 valid MHC class II exon 2 alleles, including 17 alleles of 279 bp and 6 alleles of 282 bp (Table S15). The number of alleles per individual ranged from two to nine in *Bb* and from one to nine in *Bs*, indicating that MHC class II exon 2 is a multilocus system with more than two alleles per individual. We found no significant difference in allele numbers between individuals of the two species (Wilcoxon W = 1554, p = 0.57).

We calculated the frequency of all alleles in both species (Figure 4a). We found three alleles at high frequency in *Bb,* each representing over 20% of all alleles (Bubu*01, 27.7%; Bubu*02, 20.3%; Bubu*06, 22.6%; Figure 4a; Figure S2). However, in *Bs*, we observed only two alleles exceeding 10% frequency (Bubu*14, 16.0%; Bubu*19, 15.0%; Figure 4a; Figure S2). We examined the phylogenetic relationships among the 23 alleles. All alleles were broadly distributed, exhibiting clear trans-species polymorphism between species (Figure 4b; Figure S3).

**Figure 4.**
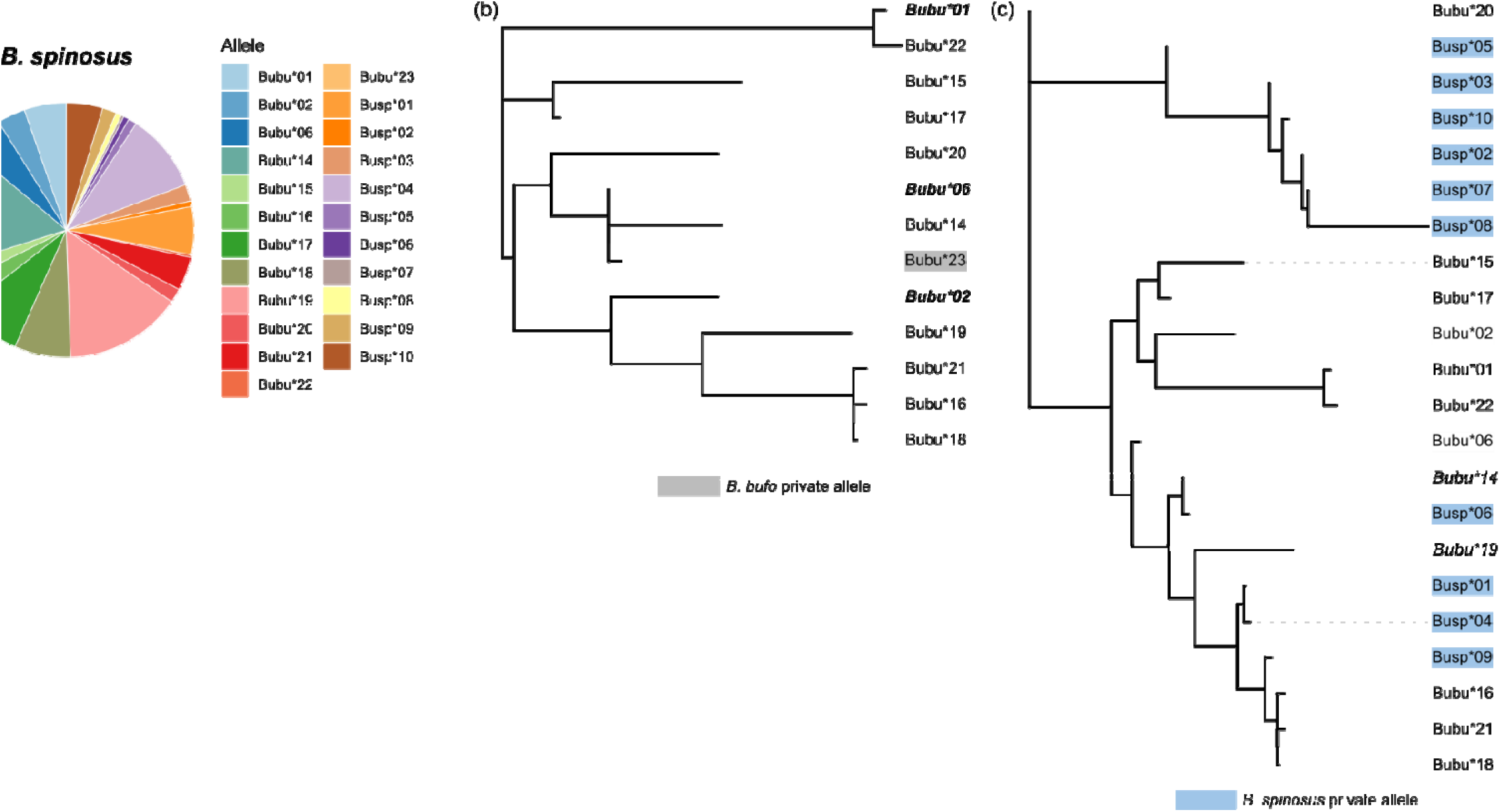
Characterization and phylogenetic relationships of MHC II exon 2 alleles. **(a)** Allelic composition and relative frequencies of MHC II exon 2 in *Bufo bufo* and *Bufo spinosus*. Pie charts illustrate the proportion of each distinct DNA allele within the respective species. **(b–c)** Maximum Likelihood (ML) phylogenetic trees depicting the evolutionary relationships of alleles identified in **(b)** *B. bufo* and **(c)** *B. spinosus*. Private alleles are highlighted in grey for *B. bufo* and blue for *B. spinosus*. The most frequent alleles (top frequencies) are indicated in bold and italics.

### MHC class II exon 2 diversity

*Bs* showed higher MHC class II exon 2 diversity compared to *Bb*, including higher Watterson’s theta (θ_w_), nucleotide diversity (π) and number of segregating sites (Table S8). We then checked allele diversity on the population level (Table S9). In *Bb*, only Stjärnamo population carried a private allele and had the greatest number of alleles and segregating sites, while Kindbäcksmåla showed highest θ_w_and π (Table S9). In *Bs*, Majada de la Vacas had the greatest number of alleles and segregating sites detected (Table S9). Population Lago Ercina carried four private alleles and showed highest θ_w_ and π among the *Bs* populations (Table S9). At the individual level, the estimated π and θ_w_ was higher in *Bb* individuals (π average = 0.0886, median = 0.09319; θ_w_ average = 0.08757, median = 0.09319) than in *Bs* individuals (π average = 0.0781, median = 0.07796; θ_w_average _=_ 0.07627, median = 0.07536). This pattern was supported by Gaussian generalized linear models (GLM; Table S16) and further confirmed by Wilcoxon rank-sum tests, which revealed significant interspecific differences for both metrics (π: W = 1954, p < 0.05; θ_w_: W = 1980, p < 0.005; Figure S4).

### 16s alpha and beta diversity

We found significant differences between non-rarefied swab samples and environmental controls in both species, confirming the effectiveness of the sampling approach (*Bb*, PERMANOVA p < 0.05, PERMDISP p < 0.05; *Bs*, PERMANOVA p < 0.05, PERMDISP p > 0.05; Figure S5-6). Skin microbiome alpha diversity was significantly higher in *Bs* than in *Bb* (Wilcoxon chao1, W = 743, p < 0.001; Wilcoxon Shannon, W = 780, p < 0.001; Wilcoxon Observed, W = 728, p < 0.001; Wilcoxon Simpson, W = 958, p < 0.01; Figure 5a). We also observed very different microbiome composition between the two species, as individuals were clustered by species on a Jaccard distance-based dendrogram (PERMANOVA, p < 0.05, Jaccard distance; Figure 5b), though the dispersion was significant (PERMDISP, p < 0.05, Jaccard distance).

**Figure 5.**
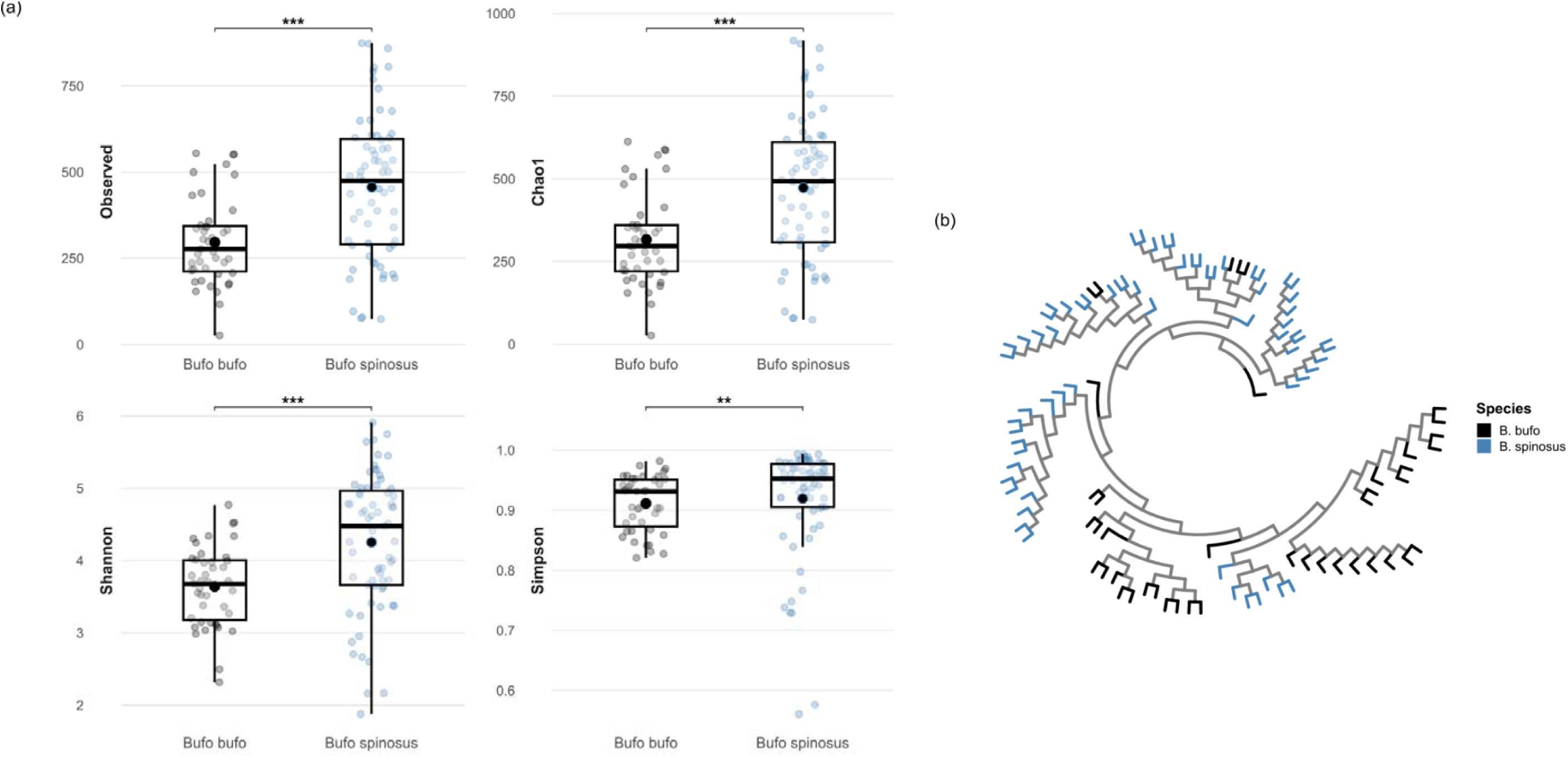
The diversity of 16S skin microbiome. **(a)** Alpha diversity, including 4 indices, Observed richness, Chao1 richness, Shannon and Simpson diversity. **(b)** beta diversity. Samples were clustered based on Jaccard distance of samples.

### Differential abundance analyses

We analyzed the differential abundance of bacterial taxa between the two species. Differential abundance analysis using LinDA identified 731 bacterial taxa significantly enriched in *Bb* and 165 taxa significantly enriched in *Bs* (log2 fold change; adjusted p < 0.05; [81]). Complementary analysis with DESeq2 revealed 159 taxa enriched in *Bb* and 208 taxa enriched in *Bs* under the same statistical thresholds (log2 fold change; adjusted p < 0.05; [82]). We selected taxa that were consistently identified as enriched by both methods, resulting in a dataset of 310 taxa, comprising 156 taxa enriched in *Bb* and 154 taxa enriched in *Bs* (Figure 6a–b; Table S17). In both species, the majority of the most enriched ASVs belonged to the phylum Pseudomonadota (Figure 6a). We observed distinct patterns of phylum-level enrichment between the two species (Figure 6b). *Bb* individuals exhibited higher relative abundances of Armatimonadota, Bacteroidota, and Pseudomonadota, whereas *Bs* individuals were enriched in Acidobacteriota, Cyanobacteriota, Deinococcota, Gemmatimonadota, Spirochaetota, Thermodesulfobacteriota, and Verrucomicrobiota (Figure 6b). To further assess the potential functional relevance of differentially abundant taxa, we compared our dataset with a published database of bacterial taxa reported to inhibit *Bd* [84]. This comparison identified 20 genera in the original dataset that have previously been associated with *Bd* inhibition (Table S18). These putative *Bd*-associated taxa were detected in both species, suggesting that both host species harbour bacterial lineages with putative antifungal potential.

**Figure 6.**
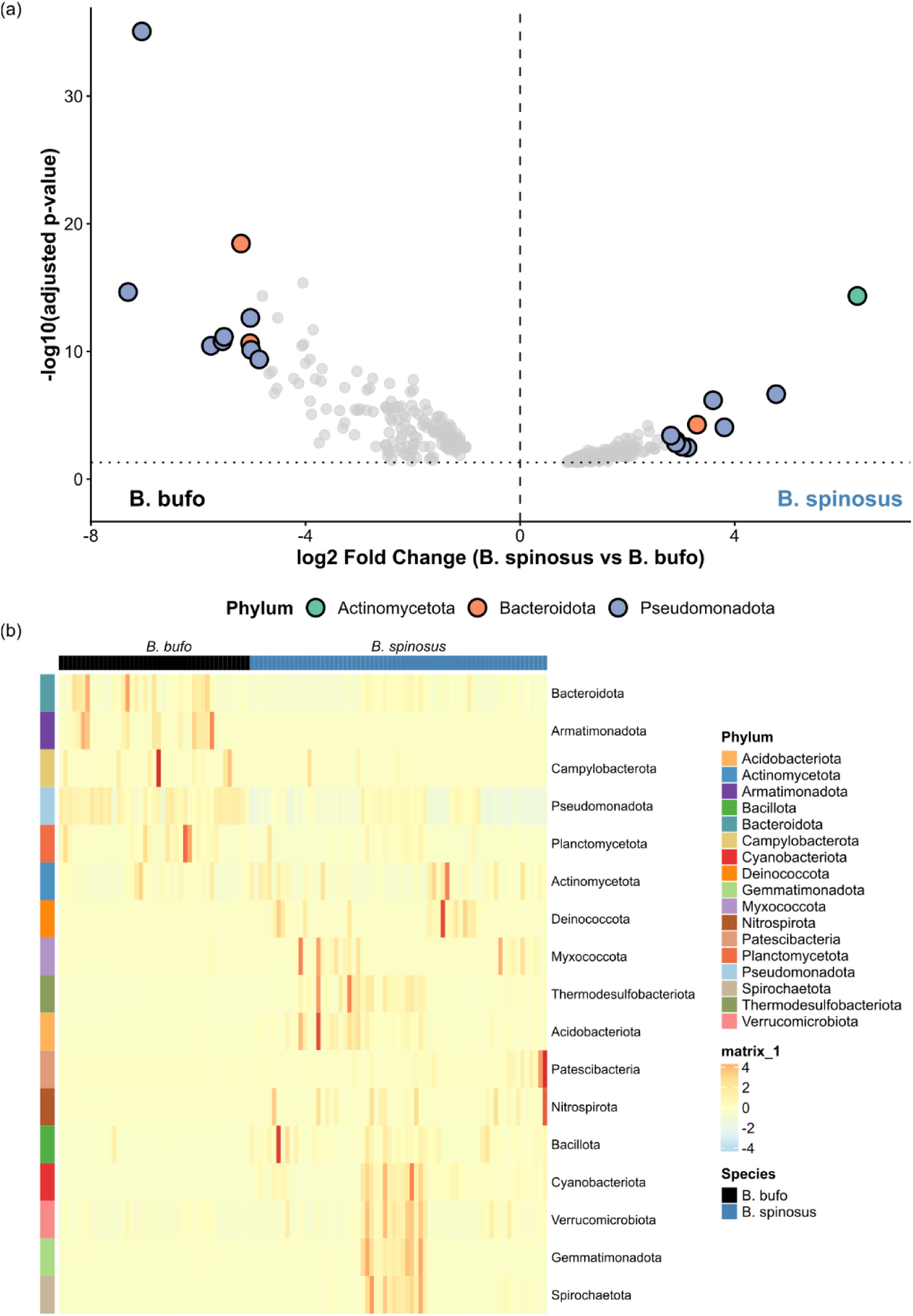
Differential skin microbiome composition between *Bufo bufo* and *B. spinosus*. **(a)** Volcano plot illustrating differentially abundant bacterial taxa between species. Each point represents a bacterial taxon, with significantly different taxa (adjusted p < 0.05) highlighted. The x-axis shows the log fold change, where positive values indicate enrichment in *B. spinosus* and negative values indicate enrichment in *B. bufo*. **(b)** Heatmap of relative abundance at the phylum level across all samples. Individuals are grouped by species as indicated by the top annotation bar (black: *B. bufo*; blue: *B. spinosus*). The warmer cell colors (red/orange) indicate higher abundance and cooler colors (white/blue) indicate lower abundance. Taxonomic classifications were shown in different colors by the left annotation bar, and the phylum names are shown on the right.

### Core microbiome alpha and beta diversity

The core taxa were shared by 70% of all individuals and it consisted of 21 bacterial taxa (Table S19). We observed contrasting patterns in alpha diversity, with higher species richness in *Bb* than in *Bs* in the indicators Chao1 and observed richness (Wilcoxon tests: Chao1, W = 2001, p < 0.05; Shannon, W = 1639, p > 0.05; observed richness, W = 2001, p < 0.05; Simpson, W = 1565, p > 0.05; Figure S7a). The core microbiomes of the two species exhibited distinct compositions (PERMANOVA, p < 0.05, Bray-Curtis distance; Figure S7b) and group dispersions (PERMDISP, p < 0.05). This divergence was dominantly driven by the differential enrichment of core taxa.

We examined the abundance of core bacterial ASVs using both LinDA and DESeq2 and identified 6 ASVs to be differentially enriched between *Bb* and *Bs* by both methods (Table S20). Four out of these six taxa found to be differentially abundant belong to the same phylum, Pseudomonadota, including three taxa of family Comamonadaceae, and one taxon of family Oxalobacteraceae.

## Discussion

We investigated genetic differences between *Bb* and *Bs*, with a focus on immune gene diversity and skin microbiome composition, to better understand the underlying mechanisms that may contribute to the potentially higher mass mortality events observed in *Bs* in the wild. While we found lower MHC class II exon 2 diversity in *Bb* compared to *Bs*, we also detected greater nucleotide divergence among alleles within individuals of *Bb* and a more diverse core microbiome in *Bb* relative to *Bs*. Together, these results suggest that greater divergence nucleotide alleles at MHC class II exon 2 and a more diverse core skin microbiome in *Bb* may contribute to the relatively lower mortality in natural populations exposed to infectious diseases. We investigated the genetic differences between *Bb* and *Bs* with a focus on immune genes to better understand the underlying differences between these two closely related species and identified differences that could potentially explain the different responses to disease. This is the first study to identify a high number of immune gene SNPs, characterize MHC class II exon 2, and profile the 16S rRNA–based skin microbiome in two closely related species that differ markedly in their infectious disease history and associated impacts.

*Bb* and *Bs* diverged following the mid–late Miocene orogenesis and Neo-Pyrenees formation (∼9.19 Mya), which isolated the ancestor of *Bs* in the Iberian Peninsula, with both species originating from a Balkan refugium that gave rise to four *Bufo* species [85]. Despite their morphological similarity [86], *Bb* underwent postglacial colonization of Scandinavia via an eastern trajectory [87], with evidence of bidirectional colonization by two genetically distinct lineages and a potential hybrid zone in central Sweden [14]. Using pcadapt to differentiate outlier from non-outlier loci and focusing on immune-related genes showing nucleotide diversity patterns consistent with balancing selection, we found distinct population genetic structure between species, with greater differentiation in *Bs* than in *Bb*. However, neutral processes such as genetic drift and geographic isolation may also contribute to this pattern [88]. The reduced differentiation in *Bb* likely reflects the geographic proximity among the sampled populations, whereas the clear divergence among *Bs* populations is consistent with their high-altitude distribution across Spain and the extensive geographic separation between sites, which is expected to limit gene flow [89,90].

Our results show reduced genetic diversity in both putatively selected loci and loci assumed to be evolving under neutrality in *Bb* compared to *Bs.* This result agrees with recent studies showing higher nuclear genetic diversity in southern Sweden than in the north, consistent with declining diversity with increasing distance from glacial refugia in *Bb* [14] and latitudinal decreases in immune gene diversity across amphibian species [11,17]. Specifically, we found that the southern species *Bs* exhibited higher overall MHC class II exon 2 diversity than *Bb*, including higher nucleotide diversity, a larger number of detected alleles, a higher number of private alleles and higher skin bacterial diversity. We also detected that several common alleles (e.g., Bubu*01, Bubu*02 and Bubu*06) occur at high frequencies in *Bb*, whereas their frequencies are substantially lower in *Bs*. Interestingly, we observed greater divergence among MHC class II alleles between individuals in *Bb* than in *Bs*, despite the overall lower MHC diversity in *Bb*, potentially reflecting differences in pathogen-mediated selection and/or demographic history between the two species [91]. Previous studies have shown that greater within-individual MHC diversity can be beneficial against pathogens by promoting immune responses to a broader range of pathogens, supported by evolutionary and geographical evidence [92,93]. Therefore, investigating immune-related genes, particularly MHC class II exon 2 diversity patterns, which have been extensively linked to *Bd* susceptibility and resistance is essential to elucidate potential associations with differing infection dynamics in these relative species.

*Bb* showed lower genetic diversity than *Bs* across three broad functional categories of immune genes. First, genes involved in complement system activation and regulation (e.g., C1S, C8B, TRIM59) showed reduced diversity in *Bb*, which is relevant for predicting infection risk given the established links between complement variability and both general disease [94,95] and *Bd* resistance [24,96,97]. Second, genes associated with immunodeficiency and immune dysfunction (CD3E, TNFRSF11A, TNFSF15) also showed lower diversity in *Bb*, where selective fixation of pathogenic variants in isolated populations could impair disease resistance [98,99]. Third, genes linked to antiviral defence and gut mucosal immunity (CD40, LTB, LIF) were similarly less diverse in *Bb;* CD40 is essential for early antiviral responses [100,101], while LTB and TNFSF15 contribute to pathogen detection and mucosal protection during infection [102–104]. Collectively, the reduced diversity across these functional categories raises concerns for Swedish *Bb* populations under potential ongoing northward pathogen spread driven by climate change. Paradoxically, *Bs* shows higher immune gene diversity yet has experienced substantial *Bd*- and *Rv*-associated mortalities (Bosch et al., 2023), suggesting that diversity alone does not confer protection and that specific variants or prolonged co-evolutionary dynamics may be driving elevated infection prevalence. Future studies quantifying the expression of these genes in experimentally infected individuals would help elucidate these potentially contrasting disease outcomes.

Additionally*, Bs* exhibited higher overall skin microbiome alpha diversity than *Bb*, and bacterial community composition differed markedly between species. Skin microbiome structure in amphibians is strongly influenced by local environmental conditions, with bacterial richness more closely associated with temperature-related variables, such as winter severity and seasonal variability, than with latitude alone [105]. Reduced microbiome diversity (dysbiosis) has been linked to increased susceptibility to infectious diseases, whereas higher diversity is often associated with lower pathogen loads and improved survival following infection [39,106]. For example, highly infected individuals can exhibit markedly reduced bacterial richness and evenness, with microbial communities dominated by a single taxonomic group [107]. Interestingly, we detected contrasting patterns of bacterial alpha diversity within the skin microbiome. Specifically, *Bb* exhibited higher richness within its core microbiome, whereas *Bs* showed higher overall skin microbiome alpha diversity. This divergence may reflect different strategies in microbiota assembly to adapt to local pathogen challenges. The higher core microbiome diversity observed in *Bb*, along with its lower *Bd* prevalence, may result from historical pathogen exposure and associated selective pressures, a pattern that has been similarly proposed to explain variation in *Bd* susceptibility across amphibian species [108]. Therefore, long-term interactions with low-virulence *Bd*-GPL strains may have favoured the persistence of a stable and diverse core microbiome that contributes to resistance. In contrast, *Bs* exhibits higher overall microbiome diversity but lower core microbiome diversity, with a substantial portion of the observed diversity represented by low-peripheral taxa that may compromise stable pathogen defense functions. Further infection experiments are needed to determine whether differences in overall and core microbiome diversity directly influence resistance to *Bd* infection.

Differential abundance analyses identified several bacterial taxa previously linked to pathogen inhibition, particularly against *Bd*. Notably, multiple genera belonging to the families Oxalobacteraceae (e.g. *Janthinobacterium*), Pseudomonadaceae (*Pseudomonas*), and Weeksellaceae (*Chryseobacterium*) were enriched in *Bb*, whereas members of the family Aeromonadaceae (*Aeromonas*) were enriched in *Bs*. Genera such as *Janthinobacterium* and *Pseudomonas* have been repeatedly shown in previous studies to include isolates capable of inhibiting *Bd* growth in vitro, primarily through the production of antifungal metabolites [109–112]. Although evidence for *Bd* inhibition by these taxa is primarily derived from in vitro assays [109–112], their presence and relative abundance in *Bb* may offer a complementary explanation for the comparatively low *Bd* infection prevalence observed in Scandinavian common toad populations. Our results suggest that interspecific differences in skin microbial diversity and community composition may be associated with both historical species divergence and subsequent local adaptation to regional environmental conditions [37,113,114]. However, more extensive spatial and temporal sampling across a wider range of locations is needed to better assess the relationship between skin microbiome composition and disease resistance, as well as to determine how microbiome shifts in captivity may influence experimentally observed mortality compared to natural populations.

## Conclusions

In summary, the variation in immune gene diversity and skin microbiome composition between *Bb* and *Bs* may be explained by species-specific demographic histories and habitat characteristics. Post-glacial colonization reduced immune gene diversity in northern *Bb*, yet adaptive tolerance and a stable, diverse core microbiome may mitigate disease impact. Conversely, *Bs* exhibits higher overall microbiome diversity—potentially reflecting elevated parasite pressures and greater encounter rates with pathogens at southern latitudes—but lower core stability driven by rare taxa, aligning with increased susceptibility. These patterns highlight how genetic variation, microbial ecology, and latitudinal parasite gradients interact to shape amphibian disease outcomes under climate-driven pathogen spread.

## Acknowledgements

We thank Ramin Eghbal, Sebastian Blom, Alex Singer and Tommy Lindström for valuable assistance in sampling in Sweden. The National Genomics Infrastructure in Stockholm funded by Science for Life Laboratory, Knut and Alice Wallenberg Foundation and the Swedish Research Council, and NAISS are acknowledged for assistance with massively parallel sequencing and access to the UPPMAX computational infrastructure

## Data accessibility

Electronic supplementary material is available online: https://figshare.com/s/f71d13f5ce8a68dac66f

BioProject and associated SRA metadata are available at https://dataview.ncbi.nlm.nih.gov/object/PRJNA1445026?reviewer=qa7kqqnu0598iimic859jg3175 for reviewing purposes. Final Bioproject number will be added before publication.

## Authors contribution

**SE**: lab work, data curation, formal analyses, writing original draft; **ZH**: lab work, data curation, formal analyses, writing original draft; **TF**: statistical-bioinformatic support, writing-review and editing; **MRP**: statistical-bioinformatic support, writing-review and editing, **NC**: field work, writing-review and editing; **BJ**: field work, writing-review and editing; **TB:** field work, writing-review and editing; **AL**: conceptualization, funding, writing-review and editing; **JH**: conceptualization, funding, writing-review and editing; **CCM**: conceptualization, funding, writing original draft, writing-review and editing.

