## Supplementary materials for "Latitudinal Patterns in Immunogenetic and Microbiome Diversity in two anuran species: *Bufo bufo* and *Bufo spinosus*"

Short title: Patterns of immunogenetic and microbiome diversity in two closely related anurans

Eleonore Susi[1]*, Zhenzhen He[1]*, Filip Thörn[2], Patrik Rödin-Mörch[1], Niki Chondrelli[1], Barbora Thumsova[3], Jaime Bosch[4], Anssi Laurila[1], Jacob Höglund[1], Maria Cortazar-Chinarro[1]

*These authors have contributed equally to the work and share first authorship

**1** Animal Ecology/Department of Ecology and Genetics, Uppsala University, Uppsala, Sweden

**2** Evolutionary Biology/Department of Ecology and Genetics, Uppsala University, Uppsala, Sweden

**3** Department of Ecology, Faculty of Science, Charles University, 128 00 Prague, Czech Republic

**4** Biodiversity Research Institute - IMIB (University of Oviedo, CSIC, Principality of Asturias), 33600 Mieres, Spain

**Corresponding authors:**

**Eleonore Susi:**

**Zhenzhen He:**

**Supplementary material**

**Additional Information 1**

MHC class II exon 2 loci were amplified using the forward primer 2F347 (5’-TGACCCTCTGCTCTCCATT-3’) and reverse primer 2R307b (5’-ATAATTCAGTATATACAGGGTCTCACC-3’), as described by Zeisset and Beebee [1]. As specified in Cortázar-Chinarro et al. [2], both forward and reverse primers were designed to include an 8 bp barcode and a sequence of three random nucleotides (NNN) to facilitate amplicon identification during sequencing. Samples were amplified separately using combinations of 6 forward and 8 reverse primers (Table S2) in reaction mixes freshly prepared according to Table S3. PCR amplification was performed using Phusion High-Fidelity DNA Polymerase (Thermo Fisher Scientific) under the following thermal cycling conditions: an initial denaturation at 97 °C for 30 s; 30 cycles of denaturation at 98 °C for 10 s, annealing at 57 °C for 20 s, and extension at 72 °C for 15 s; and a final extension at 72 °C for 8 min. Following amplification, PCR products were inspected on a 1.5% agarose gel stained with GelRed (Biotium) to estimate concentration via band intensity, and samples that failed to amplify (n = 7) were excluded. Subsequently, 7–9 samples with similar concentrations were pre-pooled and resolved on a 1.5% agarose gel for size selection. To ensure high library purity, target bands were excised and purified using the MinElute Gel Extraction Kit (Qiagen).The DNA concentration of each pre-pool was then quantified using a Qubit fluorometer (Thermo Fisher Scientific) to enable the creation of four equimolar final pools (A–D), each containing 29–30 samples. To allow pool identification during sequencing, each final pool was assigned a unique single adaptor index using the ThruPLEX DNA-Seq Kit (Takara Bio), following the manufacturer’s instructions (Table S4). Final library quality was assessed using an Agilent Bioanalyzer before sequencing at NGI, SciLifeLab (Uppsala) on the Illumina MiSeq platform.


1. Zeisset I, Beebee TJC. 2013 Bufo MHC class II loci with conserved introns flanking exon 2: cross-species amplification with common primers. Conservation Genet Resour 5, 211–213. (doi:10.1007/s12686-012-9770-y)
2. Cortázar-Chinarro M, Lattenkamp EZ, Meyer-Lucht Y, Luquet E, Laurila A, Höglund J. 2017 Drift, selection, or migration? Processes affecting genetic differentiation and variation along a latitudinal gradient in an amphibian. BMC Evol Biol 17, 189. (doi:10.1186/s12862-017-1022-z)

### Additional Information 2

In PCR1, the V4 region of the bacterial 16S rRNA gene was amplified using primers 515F (5′-GTGCCAGCMGCCGCGGTAA-3′) and 806R (5′-GGACTACHVGGGTWTCTAAT-3′), following Varg et al. [1]. The dual indices attached to both ends of the amplicons used in PCR2 were shown Table S5. Both PCRs were performed using Phusion High-Fidelity DNA Polymerase (Thermo Fisher Scientific) with reaction mixes freshly prepared for each amplification. Bovine Serum Albumin (BSA, Thermo Fisher) was included in the first PCR mix to improve amplification efficiency, whereas BSA was not used in the second PCR to avoid over-amplification (Table S6). Thermal cycling conditions for both PCRs consisted of an initial denaturation at 98 °C for 30 s, followed by denaturation at 98 °C for 10 s, annealing at 56 °C for 20 s, and extension at 72 °C for 15 s, with a final extension at 72 °C for 8 min. The number of amplification cycles was 32 for PCR1 and was reduced to 25 during PCR2 to prevent over-amplification. PCR products were purified using the Agencourt AMPure XP purification kit (Beckman Coulter Inc.) after both PCR steps. Following PCR2, amplicon concentrations were quantified using the Quant-iT PicoGreen dsDNA Assay Kit (Invitrogen Life Technologies). Finally, PCR products were balanced to equimolar concentrations and pooled, and library quality was checked using a Bioanalyzer (Agilent Technologies) before sequencing at SciLifeLab (Uppsala, Sweden) on the Illumina MiSeq platform.


1. Varg JE, Outomuro D, Kunce W, Kuehrer L, Svanbäck R, Johansson F. 2022 Microplastic exposure across trophic levels: effects on the host–microbiota of freshwater organisms. Environmental Microbiome 17, 36. (doi:10.1186/s40793-022-00429-x)

### Additional information 3. 16S rRNA metabarcoding data workflow

Forward and reverse reads were trimmed to 240 bp and 160 bp, respectively. During quality filtering, reads were truncated at the first base with a quality score ≤ 8; all other parameters were set to default. DADA2 [1] was also employed for error correction, chimera removal, and generation of amplicon sequence variants (ASVs) from 16S rRNA gene sequences, followed by taxonomic annotation of bacterial ASVs. Taxonomic classification was performed using the SILVA reference database (version 138.2) formatted for DADA2 [2]. We then employed the phyloseq package in R for subsequent data processing [3]. Contaminants were removed using both prevalence and frequency methods with default parameters in the decontam R package [4]. Taxa assigned to "Archaea", "Eukaryota", "Chloroplast", "Mitochondria", or labeled as "Unknown" were removed. We also removed singletons and eliminated ASVs with less than 10 reads across all samples [5].

### Additional information 4. MHC class II data processing

We employed the Degree of Change (DOC) method to distinguish true alleles from sequencing artefacts [1]. Variants were first ordered by sequencing depth in descending order, and the degree of change in coverage between successive variants was then calculated to identify a breakpoint in the decline [1]. This breakpoint was subsequently used to separate true alleles from artefactual sequences [1]. True alleles were aligned using MEGA 12 [2]. Alleles containing stop codons were excluded from all subsequent analyses. The remaining alleles were named following the nomenclature suggested by Klein [3], consisting of a species abbreviation followed by an asterisk and a unique allele number (e.g., Bubu*01). The allele numbering in this study was assigned in accordance with the MHC gene variants characterized by Cortazar-Chinarro et al. [4].


1. Lighten J, Van Oosterhout C, Paterson IG, McMullan M, Bentzen P. 2014 Ultra-deep Illumina sequencing accurately identifies MHC class IIb alleles and provides evidence for copy number variation in the guppy (Poecilia reticulata). Molecular Ecology Resources 14, 753–767. (doi:10.1111/1755-0998.12225)
2. Kumar S, Stecher G, Suleski M, Sanderford M, Sharma S, Tamura K. 2024 MEGA12: Molecular Evolutionary Genetic Analysis Version 12 for Adaptive and Green Computing. Molecular Biology and Evolution 41, msae263. (doi:10.1093/molbev/msae263)
3. Klein J. 1975 Biology of the Mouse Histocompatibility-2 Complex: Principles of Immunogenetics Applied to a Single System. Berlin, Heidelberg: Springer Berlin Heidelberg.
4. Cortazar-Chinarro M, Meurling S, Schroyens L, Siljestam M, Richter-Boix A, Laurila A, Höglund J. 2022 Major Histocompatibility Complex Variation and Haplotype Associated Survival in Response to Experimental Infection of Two Bd-GPL Strains Along a Latitudinal Gradient. Front. Ecol. Evol. 10, 915271. (doi:10.3389/fevo.2022.915271)

### Additional information 5. Nucleotide diversity and GO´s filtration

The nucleotide diversity was calculated per site (--site-pi), only including sites that are found in >90% individuals (--max-missing 0.9). The *B. bufo* annotations file (GCF_905171765.1; NCBI RefSeq assembly) and the GO-basic.obo file were used to extract immune-related genes that matched the root term GO:0002376 ("immune system process") or its descendants, and were marked as "involved in". If a match was not found, it was treated as a non-immune-related GO, and removed from further analyses. The file with gene annotations and GO names was combined with the nucleotide diversity dataset. The genes were divided into eight categories that would describe the immune system processes based on their GO_Name (e.g. "Adaptive Immunity", "Antigen Processing and Presentation", "Cell Death and Proliferation", "Cytokine and Interferon Response", "Immunoglobulin and Humoral Response", "Innate Immunity", "Leukocyte Migration and Chemotaxis", "Lymphoid Organ and Development", Other). GOs that did not match a category were listed as "Other" (e.g some immune genes only had the GO:0006955 - "immune response"). The genes identified as "Others" were manually searched based on the Gene_ID to try to identify their role in immunity and the corresponding category. If the gene was uncharacterized and no information was found, it would remain listed as "Other". The duplicates of genes (multiple immune-related GO_Names) matching the same category on our list were combined (only one duplicate left). Genes that matched various categories were manually searched to identify the best fit based on their function, discarding other matches.

### Additional Information 6: Raw data processing results

**Whole genome raw data processing:** The sample genomes of 30 individuals were re-sequenced with an estimated coverage depth of 14.02X (sd=1.65), with samples ranging between 10.53X and 17.07X. All samples had an average read length of 151bp and had approximately 185- 310M reads per sample. A total of 4.7 B reads per sample were aligned to the reference genome (GCF_905171765.1; NCBI RefSeq assembly) with an estimated coverage of ~70 billion bp. After final filtering, 2 972 053 SNPs passed, and 88,634 SNPs fell into regions that matched our list of immune genes (N=1102).

**MHC class II exon 2 sequencing raw data processing:** Raw MiSeq sequencing of the MHC class II exon 2 amplicon library generated approximately 21 million reads containing primers and barcodes. Samples with poor sequencing quality (n = 3) were removed, resulting in a final dataset comprising 72 *B. spinosus* and 43 *B. bufo* individuals. The average number of reads per individual was 48912, ranging from 13198 to 117504. From the 115 individuals, we identified 23 distinct MHC class II exon 2 alleles. Three of these alleles, Bubu*06, Bubu*02, and Bubu*01, were previously described by Cortazar-Chinarro [1].

**16S rRNA sequencing raw data processing:** Raw reads from the 16s metabarcoding sequencing generated 28.3 million reads of both forward and reverse direction. Following sequence assembly, quality control, and decontamination, the final dataset contained 5.1 million microbial 16S rRNA sequences and included 13993 identified taxa. We removed N=10 samples from *B. spinosus* from location Barranco Lugar due to the low sequencing depth. After removing low quality samples, we obtained a final dataset of 5.0 million reads and 13680 taxa (67 B. spinosus and 43 B. bufo) for consecutive analyses. The most abundant phyla were Pseudomonadota (61.2%), Actinomycetota (19.0%), Bacteroidota (13.1%) and Verrucomicrobiota (1.50%), while all other detected phyla each represented less than 1% of total reads (Figure S8).

1. Cortazar-Chinarro M, Meurling S, Schroyens L, Siljestam M, Richter-Boix A, Laurila A, Höglund J. 2022 Major Histocompatibility Complex Variation and Haplotype Associated Survival in Response to Experimental Infection of Two Bd-GPL Strains Along a Latitudinal Gradient. Front. Ecol. Evol. 10, 915271. (doi:10.3389/fevo.2022.915271)

**Table S1. Location details for sample collection**

| Species | Country | Location | Population | Coordinates |
|---|---|---|---|---|
| *B.spinosus* | Spain | PN Picos de Europa | Lago Ercina | 43.26740, -4.98176 |
| | | PN Sierra de Guadarrama | Laguna de los Pajaros | 40.86017, -3.94806 |
| | | | Laguna Grande | 40.83998, -3.95735 |
| | | Sierra Nevada | Majada de la vacas | 37.06584, -2.81369 |
| | | | Soportuja | 36.96326, -3.39301 |
| | | Ordesa | Barranco Lugar | 42.59640, 0.12439 |
| *B.bufo* | Sweden | Kalmar | Stjärnamo | 56.757863, 16.06023 |
| | | | Kindbäcksmåla | 56.507112, 15.754273 |
| | | | Påryd | 56.608585, 15.952671 |

**Table S2. The barcodes used in MHC II exon 2 amplicon library construction.**

| forward name | Tag | reverse name | tag |
|---|---|---|---|
| VirMiseqF1 | ACTAGATC | VirMiseqR1 | ACATGTGT |
| VirMiseqF2 | ATGATCGC | VirMiseqR2 | ACTCTGCT |
| VirMiseqF3 | CATCAGTC | VirMiseqR3 | CGTATACA |
| VirMiseqF4 | CTATGCTA | VirMiseqR4 | CTGCGTAC |
| VirMiseqF6 | TAGCTAGT | VirMiseqR5 | GATCGCGA |
| | | VirMiseqR6 | GATGATCT |
| | | VirMiseqR7 | GTCGTAGA |
| | | VirMiseqR8 | TCTACTGA |

**Table S3. PCR reaction mixture for amphibian MHC class II exon 2 amplification using barcoded primers.**

| Reagents | Stock Concentration | Volume per Reaction (μL) | Final Concentration |
|---|---|---|---|
| ddH2O | N/A | 22.1 | N/A |
| 5X HF buffer | 5X | 6 | 1X |
| dNTP | 10 mM | 0.6 | 0.20 mM |
| forward primer | 10 μM | 0.75 | 0.25 μM |
| reverse primer | 10 μM | 0.75 | 0.25 μM |
| Phusion taq | 2 U/μL | 0.3 | 0.019 U/μL |
| Template DNA | N/A | 0.3 | N/A |

**Table S4. Sample indices used in MHC II exon 2 amplicon library Miseq sequencing (ThruPlex DNA prep; TakaraBio).**

| Pool name | Sample index |
|---|---|
| A | GCCAAT |
| B | CAGATC |
| C | ACTTGA |
| D | GATCAG |

**Table S5. The barcodes used in bacterial 16S rRNA (V4 region) amplicon library construction.**

| 7i ID | 7i sequence | 5i ID | 5i sequence |
|---|---|---|---|
| 705 | GGACTCCT | 521 | CTTGCTTT |
| 706 | TAGGCATG | 522 | GGCTTCAA |
| 707 | CTCTCTAC | 523 | AATCGGCA |
| 708 | CAGAGAGG | 524 | GGTTCAAA |
| 709 | GCTACGCT | 525 | ACTTCGAC |
| 710 | CGAGGCTG | 526 | TGACTTGC |
| 711 | AAGAGGCA | 527 | TAGGACCT |
| 712 | GTAGAGGA | 528 | GGAGACTT |
| 733 | CTTATACC | | |
| 735 | TCTAGCTG | | |
| 736 | CCATAGCA | | |
| 738 | GGTATAGC | | |
| 739 | GGTTATGC | | |
| 740 | TAGGCAAG | | |
| 741 | TTGTCCAT | | |
| 743 | TCTAGGCA | | |
| 701 | TCGCCTTA | | |
| N720 | AGGCTCCG | | |
| N721 | GCAGCGTA | | |
| N722 | CTGCGCAT | | |

**Table S6. PCR reaction mixture for bacterial 16S rRNA gene (V4 region) amplification using barcoded primers.**

| | | PCR1 | | PCR2 | |
|---|---|---|---|---|---|
| Reagents | Stock concentration | Volume per Reaction (µL) | Final Concentration | Volume per Reaction (µL) | Final Concentration |
| ddH2O | N/A | 13.3 | N/A | 24.1 | N/A |
| HF buffer | 5X | 4 | 1X | 3 | 0.50X |
| dNTP | 10 mM | 0.4 | 0.20 mM | 0.6 | 0.20 mM |
| forward primer | 10 µM | 0.25 | 0.13 µM | 0.75 | 0.25 µM |
| reverse primer | 10 µM | 0.25 | 0.13 µM | 0.75 | 0.25 µM |
| Phusion taq | 2 U/µL | 0.2 | 0.020 U/µL | 0.3 | 0.020 U/µL |
| BSA | 20 mg/mL | 0.6 | 0.6 mg/mL | N/A | N/A |
| sample | N/A | 1 | N/A | 1 | N/A |

**Table S7. Variant filtering arguments used to remove low-quality reads**

| Variant filtering with GATK | Final filtering with vcftools |
|---|---|
| --filter-name "SNP_filter0" \ | --max-missing 0.9 \ |
| --filter-expression "QUAL < 30" \ | --max-alleles 2 \ |
| --filter-name "SNP_filter1" \ | --minDP 8 \ |
| --filter-expression "MQ < 40.00" \ | --maxDP 42 \ |
| --filter-name "SNP_filter2" \ | --remove-filtered-all \ |
| --filter-expression "SOR > 4.000" \ | |
| --filter-name "SNP_filter3" \ | |
| --filter-expression "QD < 2.00" \ | |
| --filter-name "SNP_filter4" \ | |
| --filter-expression "FS > 60.000" \ | |
| --filter-name "SNP_filter5" \ | |
| --filter-expression "MQRankSum < -12.5" \ | |
| --filter-name "SNP_filter6" \ | |
| --filter-expression "ReadPosRankSum < -8.000" \ | |

**Table S8. MHC class II exon diversity estimated at the species level.**

Five metrics were included: number of alleles (Ai, the number of different alleles found in the indicated species), number of private alleles (Pi, the number of alleles unique to the indicated species), Watterson's theta ($\theta_W$), nucleotide diversity ($\pi$) and number of segregating sites (S).

| Species | Number of alleles (Ai) | Number of private alleles (Pi) | Watterson's theta per site ($\theta_W$) | Nucleotide diversity ($\pi$) | Number of segregating sites (S) |
|---|---|---|---|---|---|
| *B. bufo* | 13 | 1 | 0.06815 | 0.07913 | 59 |
| *B. spinosus* | 22 | 10 | 0.06883 | 0.09383 | 70 |

**Table S9. MHC class II exon diversity estimated at the population level.**

Five metrics were included: number of alleles (Ai, the number of different alleles found in the indicated species), number of private alleles (Pi, the number of alleles unique to the indicated species), Watterson's theta ($\theta_W$), nucleotide diversity ($\pi$) and number of segregating sites (S).

| Species | Population | Allele number (Ai) | Private allele number (Pi) | Watterson's theta per site ($\theta_W$) | Nucleotide diversity ($\pi$) | Number of segregating sites (S) |
|---|---|---|---|---|---|---|
| | Stjärnamo | 12 | 1 | 0.06884 | 0.07435 | 58 |
| *B. bufo* | Kindbäcksmåla | 11 | 0 | 0.06975 | 0.08159 | 57 |
| | Påryd | 9 | 0 | 0.06858 | 0.07517 | 52 |
| | Laguna Grande | 8 | 0 | 0.07465 | 0.07668 | 54 |
| | Majada de la vacas | 17 | 0 | 0.07421 | 0.09290 | 70 |
| *B. spinosus* | Soportujar | 13 | 0 | 0.06584 | 0.07288 | 57 |
| | Laguna de los Pajaros | 10 | 0 | 0.06842 | 0.06746 | 54 |
| | Lago Ercina | 10 | 4 | 0.08489 | 0.1037 | 67 |

**Table S10. Individual-level nucleotide diversity and Watterson's theta.**

Individual nucleotide diversity (π) and Watterson's theta ($\theta_W$) per site calculated for each individual based on MHC class II exon 2 alleles. Population and species identity are indicated for each individual.

| Individual | Species | Population | Pi | ThetaW |
|---|---|---|---|---|
| X10 | *B. bufo* | Kindbäcksmåla | 0.07796 | 0.07188 |
| X11 | *B. bufo* | Kindbäcksmåla | 0.08385 | 0.07050 |
| X12 | *B. bufo* | Kindbäcksmåla | 0.07749 | 0.07754 |
| X20 | *B. bufo* | Kindbäcksmåla | 0.09319 | 0.09319 |
| X21 | *B. bufo* | Kindbäcksmåla | 0.09319 | 0.09319 |
| X32 | *B. bufo* | Kindbäcksmåla | 0.07384 | 0.07398 |
| X33 | *B. bufo* | Kindbäcksmåla | 0.09319 | 0.09319 |
| X34 | *B. bufo* | Kindbäcksmåla | 0.09319 | 0.09319 |
| X35 | *B. bufo* | Kindbäcksmåla | 0.09319 | 0.09319 |
| X36 | *B. bufo* | Kindbäcksmåla | 0.09319 | 0.09319 |
| X37 | *B. bufo* | Kindbäcksmåla | 0.09319 | 0.09319 |
| X6 | *B. bufo* | Kindbäcksmåla | 0.08158 | 0.07607 |
| X7 | *B. bufo* | Kindbäcksmåla | 0.07431 | 0.07064 |
| X8 | *B. bufo* | Kindbäcksmåla | 0.07749 | 0.07754 |
| X9 | *B. bufo* | Kindbäcksmåla | 0.07168 | 0.07168 |
| B8289 | *B. spinosus* | Lago Ercina | 0.11111 | 0.11111 |
| B8290 | *B. spinosus* | Lago Ercina | 0.10155 | 0.09189 |
| B8291 | *B. spinosus* | Lago Ercina | 0.06810 | 0.06810 |
| B8292 | *B. spinosus* | Lago Ercina | 0.06810 | 0.06810 |
| B8293 | *B. spinosus* | Lago Ercina | 0.12903 | 0.12903 |
| B8294 | *B. spinosus* | Lago Ercina | 0.06810 | 0.06810 |
| B8305 | *B. spinosus* | Lago Ercina | 0.06810 | 0.06810 |
| B8306 | *B. spinosus* | Lago Ercina | 0.06810 | 0.06810 |
| B8307 | *B. spinosus* | Lago Ercina | 0.06452 | 0.06452 |
| B8309 | *B. spinosus* | Lago Ercina | 0.06810 | 0.06810 |
| B8310 | *B. spinosus* | Lago Ercina | 0.11231 | 0.10992 |
| B8312 | *B. spinosus* | Lago Ercina | 0.08961 | 0.08602 |
| B8313 | *B. spinosus* | Lago Ercina | 0.11470 | 0.11231 |
| B8314 | *B. spinosus* | Lago Ercina | 0.06810 | 0.06810 |
| B8325 | *B. spinosus* | Lago Ercina | 0.15054 | 0.15054 |
| B8326 | *B. spinosus* | Lago Ercina | 0.10753 | 0.10514 |
| B8327 | *B. spinosus* | Lago Ercina | 0.11470 | 0.11231 |
| B8328 | *B. spinosus* | Lago Ercina | 0.11231 | 0.10992 |
| X117 | *B. spinosus* | Laguna de los Pajaros | 0.06213 | 0.06213 |
| X118 | *B. spinosus* | Laguna de los Pajaros | 0.05018 | 0.05018 |
| X119 | *B. spinosus* | Laguna de los Pajaros | 0.04946 | 0.04817 |
| X120 | *B. spinosus* | Laguna de los Pajaros | 0.05018 | 0.05018 |
| X121 | *B. spinosus* | Laguna de los Pajaros | 0.05018 | 0.05018 |
| X122 | *B. spinosus* | Laguna de los Pajaros | 0.08363 | 0.08320 |

| | | | | |
|---|---|---|---|---|
| X123 | *B. spinosus* | Laguna de los Pajaros | 0.03345 | 0.03345 |
| X124 | *B. spinosus* | Laguna de los Pajaros | 0.13620 | 0.13620 |
| X125 | *B. spinosus* | Laguna de los Pajaros | 0.11470 | 0.10992 |
| X126 | *B. spinosus* | Laguna de los Pajaros | 0.04062 | 0.04062 |
| X127 | *B. spinosus* | Laguna de los Pajaros | 0.06093 | 0.05494 |
| X101 | *B. spinosus* | Laguna de los Pajaros | 0.05018 | 0.05018 |
| X102 | *B. spinosus* | Laguna de los Pajaros | 0.10992 | 0.10753 |
| X103 | *B. spinosus* | Laguna de los Pajaros | 0.09916 | 0.09775 |
| X104 | *B. spinosus* | Laguna de los Pajaros | 0.06093 | 0.05494 |
| X106 | *B. spinosus* | Laguna de los Pajaros | 0.08961 | 0.08946 |
| X107 | *B. spinosus* | Laguna de los Pajaros | 0.08674 | 0.08602 |
| X108 | *B. spinosus* | Laguna de los Pajaros | 0.06093 | 0.05494 |
| X109 | *B. spinosus* | Laguna de los Pajaros | 0.12545 | 0.12545 |
| X110 | *B. spinosus* | Laguna de los Pajaros | 0.10096 | 0.09971 |
| B8357 | *B. spinosus* | Majada de la vacas | 0.05615 | 0.05474 |
| B8358 | *B. spinosus* | Majada de la vacas | 0.08022 | 0.07607 |
| B8359 | *B. spinosus* | Majada de la vacas | 0.05376 | 0.05376 |
| B8360 | *B. spinosus* | Majada de la vacas | 0.09379 | 0.09189 |
| B8361 | *B. spinosus* | Majada de la vacas | 0.09498 | 0.08993 |
| B8362 | *B. spinosus* | Majada de la vacas | 0.06523 | 0.06022 |
| B8363 | *B. spinosus* | Majada de la vacas | 0.00358 | 0.00358 |
| B8364 | *B. spinosus* | Majada de la vacas | 0.10992 | 0.10753 |
| B8365 | *B. spinosus* | Majada de la vacas | 0.09319 | 0.09319 |
| B8366 | *B. spinosus* | Majada de la vacas | 0.06523 | 0.06022 |
| B8367 | *B. spinosus* | Majada de la vacas | 0.04659 | 0.04659 |
| B8368 | *B. spinosus* | Majada de la vacas | 0.09976 | 0.09971 |
| B8369 | *B. spinosus* | Majada de la vacas | 0.06998 | 0.06583 |
| B8370 | *B. spinosus* | Majada de la vacas | 0.04719 | 0.04301 |
| B8371 | *B. spinosus* | Majada de la vacas | 0.08602 | 0.08602 |
| X13 | *B. bufo* | Påryd | 0.08602 | 0.08602 |
| X14 | *B. bufo* | Påryd | 0.07937 | 0.07607 |
| X15 | *B. bufo* | Påryd | 0.08662 | 0.08602 |
| X16 | *B. bufo* | Påryd | 0.09319 | 0.09319 |
| X17 | *B. bufo* | Påryd | 0.10753 | 0.10753 |
| X23 | *B. bufo* | Påryd | 0.07945 | 0.08211 |
| X38 | *B. bufo* | Påryd | 0.09319 | 0.09319 |
| X39 | *B. bufo* | Påryd | 0.09319 | 0.09319 |
| X40 | *B. bufo* | Påryd | 0.10036 | 0.10036 |
| X41 | *B. bufo* | Påryd | 0.09319 | 0.09319 |
| X42 | *B. bufo* | Påryd | 0.09319 | 0.09319 |
| X43 | *B. bufo* | Påryd | 0.09319 | 0.09319 |
| X44 | *B. bufo* | Påryd | 0.10036 | 0.10036 |
| X111 | *B. spinosus* | Soportujar | 0.07578 | 0.07050 |

| | | | | |
|---|---|---|---|---|
| X112 | *B. spinosus* | Soportujar | 0.07616 | 0.06989 |
| X113 | *B. spinosus* | Soportujar | 0.07527 | 0.07742 |
| X128 | *B. spinosus* | Soportujar | 0.08554 | 0.08163 |
| X129 | *B. spinosus* | Soportujar | 0.06571 | 0.06452 |
| X130 | *B. spinosus* | Soportujar | 0.08136 | 0.08258 |
| X131 | *B. spinosus* | Soportujar | 0.08602 | 0.08602 |
| X132 | *B. spinosus* | Soportujar | 0.08578 | 0.08163 |
| X133 | *B. spinosus* | Soportujar | 0.07851 | 0.07607 |
| X134 | *B. spinosus* | Soportujar | 0.10394 | 0.10275 |
| X135 | *B. spinosus* | Soportujar | 0.07796 | 0.07188 |
| X136 | *B. spinosus* | Soportujar | 0.08196 | 0.08163 |
| X137 | *B. spinosus* | Soportujar | 0.08602 | 0.08602 |
| X138 | *B. spinosus* | Soportujar | 0.07796 | 0.07188 |
| X139 | *B. spinosus* | Soportujar | 0.07770 | 0.07465 |
| X1 | *B. bufo* | Stjärnamo | 0.09319 | 0.09319 |
| X18 | *B. bufo* | Stjärnamo | 0.09319 | 0.09319 |
| X19 | *B. bufo* | Stjärnamo | 0.06930 | 0.06930 |
| X2 | *B. bufo* | Stjärnamo | 0.08158 | 0.07607 |
| X22 | *B. bufo* | Stjärnamo | 0.09319 | 0.09319 |
| X25 | *B. bufo* | Stjärnamo | 0.09319 | 0.09319 |
| X26 | *B. bufo* | Stjärnamo | 0.09140 | 0.08798 |
| X27 | *B. bufo* | Stjärnamo | 0.09319 | 0.09319 |
| X28 | *B. bufo* | Stjärnamo | 0.09319 | 0.09319 |
| X29 | *B. bufo* | Stjärnamo | 0.09319 | 0.09319 |
| X3 | *B. bufo* | Stjärnamo | 0.07510 | 0.06876 |
| X30 | *B. bufo* | Stjärnamo | 0.09319 | 0.09319 |
| X31 | *B. bufo* | Stjärnamo | 0.09319 | 0.09319 |
| X4 | *B. bufo* | Stjärnamo | 0.07749 | 0.07754 |
| X5 | *B. bufo* | Stjärnamo | 0.10753 | 0.10753 |

**Table S11. Innate immunity genes with mean nucleotide diversity in *B. bufo* and *B. spinosus***

| GeneID | *B. bufo* mean_PI | *B. spinosus* mean_PI | Diff. | Abs_diff. |
|---|---|---|---|---|
| C1S | 0.106 | 0.516 | 0.410 | 0.41 |
| LOC120977404 | 0.000 | 0.405 | 0.405 | 0.4 |
| TRIM59 | 0.036 | 0.429 | 0.393 | 0.39 |
| C8B | 0.000 | 0.370 | 0.370 | 0.37 |
| LOC120986786 | 0.060 | 0.397 | 0.337 | 0.34 |
| LOC120988975 | 0.145 | 0.476 | 0.331 | 0.33 |
| LOC120977484 | 0.030 | 0.354 | 0.323 | 0.32 |
| LOC120977492 | 0.063 | 0.333 | 0.270 | 0.27 |
| LOC120997021 | 0.080 | 0.349 | 0.270 | 0.27 |
| LOC120997025 | 0.080 | 0.349 | 0.270 | 0.27 |
| LOC120977496 | 0.009 | 0.260 | 0.251 | 0.25 |
| LOC120988973 | 0.007 | 0.254 | 0.247 | 0.25 |
| LOC120978194 | 0.029 | 0.272 | 0.243 | 0.24 |
| LOC120978195 | 0.021 | 0.260 | 0.239 | 0.24 |
| TRIM62 | 0.000 | 0.238 | 0.238 | 0.24 |
| LOC120977483 | 0.055 | 0.289 | 0.234 | 0.23 |
| IRF9 | 0.008 | 0.232 | 0.224 | 0.22 |
| LOC121003259 | 0.057 | 0.279 | 0.222 | 0.22 |
| LOC120978196 | 0.000 | 0.207 | 0.207 | 0.21 |
| LOC120991695 | 0.069 | 0.276 | 0.207 | 0.21 |
| LOC121003376 | 0.043 | 0.254 | 0.211 | 0.21 |
| IRF6 | 0.000 | 0.198 | 0.198 | 0.2 |
| LOC120978140 | 0.000 | 0.202 | 0.202 | 0.2 |
| LOC120988977 | 0.000 | 0.191 | 0.191 | 0.19 |
| LOC120989234 | 0.119 | 0.289 | 0.170 | 0.17 |
| LOC121001458 | 0.163 | 0.325 | 0.163 | 0.16 |
| IRF7 | 0.150 | 0.000 | -0.150 | 0.15 |
| TICAM1 | 0.000 | 0.152 | 0.152 | 0.15 |
| C1R | 0.000 | 0.138 | 0.138 | 0.14 |
| IRF3 | 0.000 | 0.132 | 0.132 | 0.13 |
| CXCL14 | 0.000 | 0.116 | 0.116 | 0.12 |
| LOC121008994 | 0.000 | 0.121 | 0.121 | 0.12 |
| NFKB2 | 0.010 | 0.125 | 0.115 | 0.12 |
| LOC120978145 | 0.371 | 0.264 | -0.108 | 0.11 |
| MYD88 | 0.053 | 0.166 | 0.113 | 0.11 |
| LOC120988974 | 0.105 | 0.210 | 0.105 | 0.1 |
| LOC120995636 | 0.131 | 0.232 | 0.101 | 0.1 |
| LY96 | 0.133 | 0.232 | 0.099 | 0.1 |
| MASP2 | 0.149 | 0.251 | 0.102 | 0.1 |
| STING1 | 0.051 | 0.150 | 0.099 | 0.1 |
| IFIH1 | 0.103 | 0.189 | 0.086 | 0.09 |
| CMKLR1 | 0.185 | 0.265 | 0.080 | 0.08 |

| | | | | |
|---|---|---|---|---|
| LOC121007991 | 0.200 | 0.283 | 0.083 | 0.08 |
| C6 | 0.214 | 0.282 | 0.068 | 0.07 |
| MAPKAPK2 | 0.101 | 0.167 | 0.066 | 0.07 |
| TICAM2 | 0.000 | 0.066 | 0.066 | 0.07 |
| LOC120978137 | 0.221 | 0.156 | -0.065 | 0.06 |
| LOC120978139 | 0.216 | 0.271 | 0.056 | 0.06 |
| LOC120982344 | 0.170 | 0.230 | 0.060 | 0.06 |
| MASP1 | 0.100 | 0.155 | 0.055 | 0.06 |
| IRF8 | 0.043 | 0.000 | -0.043 | 0.04 |
| LOC120977489 | 0.139 | 0.175 | 0.036 | 0.04 |
| PTX3 | 0.053 | 0.093 | 0.040 | 0.04 |
| TRAF3 | 0.251 | 0.213 | -0.038 | 0.04 |
| IRAK1BP1 | 0.007 | 0.037 | 0.030 | 0.03 |
| LOC120978141 | 0.124 | 0.093 | -0.031 | 0.03 |
| LY86 | 0.124 | 0.156 | 0.033 | 0.03 |
| IRF4 | 0.075 | 0.055 | -0.020 | 0.02 |
| LOC120986788 | 0.310 | 0.327 | 0.017 | 0.02 |
| LOC120991007 | 0.079 | 0.101 | 0.021 | 0.02 |
| LOC121007567 | 0.195 | 0.216 | 0.020 | 0.02 |
| C7 | 0.130 | 0.144 | 0.014 | 0.01 |
| C8A | 0.145 | 0.138 | -0.007 | 0.01 |
| LOC120977034 | 0.128 | 0.137 | 0.010 | 0.01 |
| LOC120986787 | 0.172 | 0.162 | -0.010 | 0.01 |
| IRF1 | 0.000 | 0.000 | 0.000 | 0 |
| LOC121005258 | 0.000 | 0.000 | 0.000 | 0 |
| TLR4 | 0.000 | 0.000 | 0.000 | 0 |
| C3AR1 | 0.000 | NA | NA | NA |
| IFNL3 | 0.172 | NA | NA | NA |
| LEAP2 | 0.254 | NA | NA | NA |
| LOC120977027 | 0.000 | NA | NA | NA |
| LOC120978144 | 0.080 | NA | NA | NA |
| LOC120991669 | 0.000 | NA | NA | NA |
| LOC120991683 | 0.000 | NA | NA | NA |
| LOC120986785 | NA | 0.138 | NA | NA |

**Table S12. Adaptive immunity genes with mean nucleotide diversity in *B. bufo* and *B. spinosus***

| GeneID | *B. bufo* mean_PI | *B. spinosus* mean_PI | Diff. | Abs_diff. |
|---|---|---|---|---|
| CD40 | 0.052 | 0.422 | 0.371 | 0.37 |
| LOC121008793 | 0.031 | 0.367 | 0.335 | 0.34 |
| LOC120989200 | 0.033 | 0.339 | 0.307 | 0.31 |
| CD79B | 0.005 | 0.309 | 0.304 | 0.3 |
| CD79A | 0.000 | 0.235 | 0.235 | 0.24 |
| HHLA2 | 0.005 | 0.206 | 0.201 | 0.2 |
| CD40LG | 0.027 | 0.203 | 0.176 | 0.18 |
| LIME1 | 0.099 | 0.280 | 0.181 | 0.18 |
| BCL10 | 0.110 | 0.258 | 0.148 | 0.15 |
| TNFRSF21 | 0.144 | 0.289 | 0.145 | 0.14 |
| CD80 | 0.000 | 0.129 | 0.129 | 0.13 |
| LOC120989201 | 0.067 | 0.193 | 0.125 | 0.13 |
| TNFSF8 | 0.066 | 0.200 | 0.134 | 0.13 |
| LOC120993187 | 0.214 | 0.334 | 0.120 | 0.12 |
| LOC121007987 | 0.006 | 0.127 | 0.120 | 0.12 |
| HSPD1 | 0.037 | 0.119 | 0.082 | 0.08 |
| LAG3 | 0.014 | 0.069 | 0.055 | 0.06 |
| RAG2 | NA | 0.000 | NA | NA |

**Table S13. Lymphoid & Organ Development genes with mean nucleotide diversity in *B. bufo* and *B. spinosus***

| GeneID | *B. bufo* mean_PI | *B. spinosus* mean_PI | Diff. | Abs_diff. |
|---|---|---|---|---|
| LTB | 0.000 | 0.364 | 0.364 | 0.36 |
| CD3E | 0.000 | 0.263 | 0.263 | 0.26 |
| TNFRSF11A | 0.034 | 0.232 | 0.198 | 0.2 |
| CXCR5 | 0.066 | 0.247 | 0.182 | 0.18 |
| LTBR | 0.069 | 0.246 | 0.177 | 0.18 |
| LOC120996009 | 0.138 | 0.069 | -0.069 | 0.07 |

**Table S14. Cytokine and Interferon Response genes with mean nucleotide diversity in *B. bufo* and *B. spinosus***

| GeneID | *B. bufo* mean_PI | *B. spinosus* mean_PI | Diff. | Abs_diff. |
|---|---|---|---|---|
| LIF | 0.000 | 0.423 | 0.423 | 0.42 |
| TNFSF15 | 0.030 | 0.438 | 0.409 | 0.41 |
| IL10 | 0.055 | 0.217 | 0.162 | 0.16 |
| IL6 | 0 | NA | NA | NA |

**Table S15. Distribution of detected alleles between *Bufo spinosus* and *Bufo bufo*.**

| | *B. spinosus* | *B. bufo* |
|---|---|---|
| Unique Alleles | 10 alleles (6 alleles of 282 bp: Busp*02-03, Busp*05, Busp*07-08, Busp*10; 4 alleles of 279 bp: Busp*01, Busp*04, Busp*06, Busp*09) | 1 allele (Bubu*23) |
| Shared alleles | 12 alleles (Bubu*01, Bubu*02, Bubu06, Bubu*14-22) | |
| Total alleles | 22 alleles | 13 alleles |

**Table S16. Results of a Gaussian generalized linear model (GLM) assessing differences in individual-level nucleotide diversity (π) and Watterson's theta ($\theta_w$) between *Bufo bufo* and *Bufo spinosus*.**

Model coefficients (β), standard errors (SE), test statistics (t values), and associated p-values are shown. Species identity was included as a categorical predictor, with *B. bufo* used as the reference level.

| Model Target | Parameter | Estimate (β) | Std. Error | t value | p-value |
|---|---|---|---|---|---|
| Model 1: Nucleotide diversity (π) | Intercept (Base: *B. bufo*) | 0.0886 | 0.0032 | 27.2686 | 0 |
| | Species: *B. spinosus* | -0.008 | 0.0042 | -1.9283 | 0.0564 |
| Model 2: Watterson's theta ($\theta_w$) per site | Intercept (Base: *B. bufo*) | 0.0876 | 0.0033 | 26.7233 | 0 |
| | Species: *B. spinosus* | -0.0088 | 0.0042 | -2.1042 | 0.0377 |

**Table S17. Bacterial taxa identified as differentially abundant between *Bufo bufo* and *B. spinosus* by both LinDA and DESeq2.**

| Phylum | Genus | Enriched_Species |
|---|---|---|
| Pseudomonadota | *Rhodoferax* | *B. bufo* |
| Pseudomonadota | *Sphingomonas* | *B. bufo* |
| Bacteroidota | *Fluviicola* | *B. bufo* |
| Pseudomonadota | *Leptothrix* | *B. bufo* |
| Actinomycetota | *hgcI clade* | *B. bufo* |
| Pseudomonadota | *Aeromonas* | *B. bufo* |
| Pseudomonadota | *Polymorphobacter* | *B. bufo* |
| Campylobacterota | *Arcobacter* | *B. bufo* |
| Pseudomonadota | *Herminiimonas* | *B. bufo* |
| Pseudomonadota | *Iodobacter* | *B. bufo* |
| Actinomycetota | *hgcI clade* | *B. bufo* |
| Pseudomonadota | *Hyphomicrobium* | *B. bufo* |
| Actinomycetota | *Candidatus Planktoluna* | *B. bufo* |
| Pseudomonadota | *Novosphingobium* | *B. bufo* |
| Bacteroidota | *Ferruginibacter* | *B. bufo* |
| Pseudomonadota | *Polynucleobacter* | *B. bufo* |
| Verrucomicrobiota | *Luteolibacter* | *B. bufo* |
| Pseudomonadota | *Rhizobium* | *B. bufo* |
| Pseudomonadota | *Deefgea* | *B. bufo* |
| Pseudomonadota | *Rhodoferax* | *B. bufo* |
| Pseudomonadota | *Undibacterium* | *B. bufo* |
| Verrucomicrobiota | *Luteolibacter* | *B. bufo* |
| Pseudomonadota | *Ellin6067* | *B. bufo* |
| Pseudomonadota | *Sphingomonas* | *B. bufo* |
| Verrucomicrobiota | *Cerasicoccus* | *B. bufo* |
| Bacteroidota | *Flavobacterium* | *B. bufo* |

| | | |
|---|---|---|
| Actinomycetota | *Mycobacterium* | *B. bufo* |
| Actinomycetota | *Micrococcus* | *B. bufo* |
| Pseudomonadota | *Undibacterium* | *B. bufo* |
| Pseudomonadota | *alphaI cluster* | *B. bufo* |
| Bacteroidota | *Flavobacterium* | *B. bufo* |
| Pseudomonadota | *Undibacterium* | *B. bufo* |
| Pseudomonadota | *Saccharedens* | *B. bufo* |
| Pseudomonadota | *Roseomonas* | *B. bufo* |
| Pseudomonadota | *Massilia* | *B. bufo* |
| Pseudomonadota | *Methylorosula* | *B. bufo* |
| Pseudomonadota | *Crenothrix* | *B. bufo* |
| Verrucomicrobiota | *Chthoniobacter* | *B. bufo* |
| Pseudomonadota | *Variovorax* | *B. bufo* |
| Pseudomonadota | *alphaI cluster* | *B. bufo* |
| Pseudomonadota | *Undibacterium* | *B. bufo* |
| Pseudomonadota | *Rhodovarius* | *B. bufo* |
| Pseudomonadota | *Paucibacter* | *B. bufo* |
| Pseudomonadota | *Dechloromonas* | *B. bufo* |
| Bacteroidota | *Flavobacterium* | *B. bufo* |
| Pseudomonadota | *Iodobacter* | *B. bufo* |
| Pseudomonadota | *Rugamonas* | *B. bufo* |
| Pseudomonadota | *Rhodanobacter* | *B. bufo* |
| Pseudomonadota | *Achromatium* | *B. bufo* |
| Pseudomonadota | *Rhizorhapis* | *B. bufo* |
| Pseudomonadota | *Undibacterium* | *B. bufo* |
| Pseudomonadota | *Hyphomicrobium* | *B. bufo* |
| Pseudomonadota | *Burkholderia-Caballeronia-Paraburkholderia* | *B. bufo* |
| Pseudomonadota | *Parablastomonas* | *B. bufo* |

| | | |
|---|---|---|
| Pseudomonadota | *Undibacterium* | *B. bufo* |
| Pseudomonadota | *Rhodoferax* | *B. bufo* |
| Bacteroidota | *Fibrella* | *B. bufo* |
| Bacteroidota | *Pedobacter* | *B. bufo* |
| Pseudomonadota | *Methylotenera* | *B. bufo* |
| Pseudomonadota | *Rhodoferax* | *B. bufo* |
| Pseudomonadota | *Sphaerotilus* | *B. bufo* |
| Bacteroidota | *Chryseotalea* | *B. bufo* |
| Pseudomonadota | *Rhodanobacter* | *B. bufo* |
| Pseudomonadota | *Cereibacter* | *B. bufo* |
| Pseudomonadota | *Variovorax* | *B. bufo* |
| Actinomycetota | *Candidatus Planktophila* | *B. bufo* |
| Pseudomonadota | *Leptothrix* | *B. bufo* |
| Verrucomicrobiota | *Cephaloticoccus* | *B. bufo* |
| Pseudomonadota | *Rhodoferax* | *B. bufo* |
| Pseudomonadota | *Rhizorhapis* | *B. bufo* |
| Pseudomonadota | *Lysobacter* | *B. bufo* |
| Pseudomonadota | *Caulobacter* | *B. bufo* |
| Bacteroidota | *Flavobacterium* | *B. bufo* |
| Pseudomonadota | *Novosphingobium* | *B. bufo* |
| Pseudomonadota | *Asticcacaulis* | *B. bufo* |
| Pseudomonadota | *Candidatus Methylopumilus* | *B. bufo* |
| Pseudomonadota | *Undibacterium* | *B. bufo* |
| Pseudomonadota | *Methylocella* | *B. bufo* |
| Pseudomonadota | *Sphaerotilus* | *B. bufo* |
| Pseudomonadota | *Ferribacterium* | *B. bufo* |
| Bacteroidota | *Flavobacterium* | *B. bufo* |
| Pseudomonadota | *Rhodoferax* | *B. bufo* |
| Gemmatimonadota | *Gemmatimonas* | *B. bufo* |

| | | |
|---|---|---|
| Bacteroidota | *Fluviicola* | *B. bufo* |
| Pseudomonadota | *Phreatobacter* | *B. bufo* |
| Bacteroidota | *Flavobacterium* | *B. bufo* |
| Pseudomonadota | *Rhodoferax* | *B. bufo* |
| Bacteroidota | *Flavobacterium* | *B. bufo* |
| Bacteroidota | *Emticicia* | *B. bufo* |
| Pseudomonadota | *Polynucleobacter* | *B. bufo* |
| Pseudomonadota | *Leptothrix* | *B. bufo* |
| Actinomycetota | *hgcI clade* | *B. bufo* |
| Planctomycetota | *Pirellula* | *B. bufo* |
| Pseudomonadota | *Pseudomonas* | *B. bufo* |
| Pseudomonadota | *Candidatus Methylopumilus* | *B. bufo* |
| Pseudomonadota | *Janthinobacterium* | *B. bufo* |
| Pseudomonadota | *Caulobacter* | *B. bufo* |
| Pseudomonadota | *Dechloromonas* | *B. bufo* |
| Pseudomonadota | *Hyphomicrobium* | *B. bufo* |
| Bacteroidota | *Fluviicola* | *B. bufo* |
| Bacteroidota | *Fluviicola* | *B. bufo* |
| Pseudomonadota | *Paucibacter* | *B. bufo* |
| Pseudomonadota | *Cereibacter* | *B. bufo* |
| Bacteroidota | *Flavobacterium* | *B. bufo* |
| Bacteroidota | *Dinghuibacter* | *B. bufo* |
| Pseudomonadota | *Limnohabitans* | *B. bufo* |
| Pseudomonadota | *Undibacterium* | *B. bufo* |
| Pseudomonadota | *Duganella* | *B. bufo* |
| Pseudomonadota | *Acidovorax* | *B. bufo* |
| Pseudomonadota | *Limnohabitans* | *B. bufo* |
| Planctomycetota | *Phycisphaera* | *B. bufo* |
| Pseudomonadota | *Novosphingobium* | *B. bufo* |

| | | |
|---|---|---|
| Pseudomonadota | *Parvibium* | *B. bufo* |
| Pseudomonadota | *Asticcacaulis* | *B. bufo* |
| Actinomycetota | *Flexivirga* | *B. bufo* |
| Pseudomonadota | *Brevundimonas* | *B. bufo* |
| Pseudomonadota | *Polaromonas* | *B. bufo* |
| Pseudomonadota | *Rhodoferax* | *B. bufo* |
| Pseudomonadota | *Simplicispira* | *B. bufo* |
| Pseudomonadota | *Phreatobacter* | *B. bufo* |
| Pseudomonadota | *Ottowia* | *B. bufo* |
| Bacteroidota | *Chryseobacterium* | *B. bufo* |
| Pseudomonadota | *Hirschia* | *B. bufo* |
| Pseudomonadota | *Iodobacter* | *B. bufo* |
| Pseudomonadota | *Rhodoferax* | *B. bufo* |
| Pseudomonadota | *Luteibacter* | *B. bufo* |
| Pseudomonadota | *Herminiimonas* | *B. bufo* |
| Pseudomonadota | *UKL13-1* | *B. bufo* |
| Bacteroidota | *Flavobacterium* | *B. bufo* |
| Bacteroidota | *Sediminibacterium* | *B. bufo* |
| Pseudomonadota | *Actimicrobium* | *B. bufo* |
| Pseudomonadota | *Parablastomonas* | *B. bufo* |
| Bacteroidota | *Flavobacterium* | *B. bufo* |
| Bacteroidota | *Sediminibacterium* | *B. bufo* |
| Pseudomonadota | *Limnohabitans* | *B. bufo* |
| Pseudomonadota | *Ferribacterium* | *B. bufo* |
| Pseudomonadota | *Methylotenera* | *B. bufo* |
| Pseudomonadota | *Paucibacter* | *B. bufo* |
| Pseudomonadota | *Lichenibacterium* | *B. bufo* |
| Actinomycetota | *hgcI clade* | *B. bufo* |
| Pseudomonadota | *Georgfuchsia* | *B. bufo* |

| | | |
|---|---|---|
| Bacteroidota | *Chryseobacterium* | *B. bufo* |
| Pseudomonadota | *Methylotenera* | *B. bufo* |
| Armatimonadota | *Armatimonas* | *B. bufo* |
| Pseudomonadota | *Cypionkella* | *B. bufo* |
| Bacteroidota | *Fluviicola* | *B. bufo* |
| Pseudomonadota | *Caulobacter* | *B. bufo* |
| Actinomycetota | *hgcI clade* | *B. bufo* |
| Pseudomonadota | *Caulobacter* | *B. bufo* |
| Pseudomonadota | *Methylotenera* | *B. bufo* |
| Bacteroidota | *Flavobacterium* | *B. bufo* |
| Bacteroidota | *Lacihabitans* | *B. bufo* |
| Bacteroidota | *Chryseotalea* | *B. bufo* |
| Actinomycetota | *Candidatus Planktophila* | *B. bufo* |
| Pseudomonadota | *Rhodoferax* | *B. bufo* |
| Bacteroidota | *Flavobacterium* | *B. bufo* |
| Bacteroidota | *Runella* | *B. spinosus* |
| Bacteroidota | *Flavobacterium* | *B. spinosus* |
| Bacteroidota | *BSV13* | *B. spinosus* |
| Acidobacteriota | *Paludibaculum* | *B. spinosus* |
| Verrucomicrobiota | *ADurb.Bin063-1* | *B. spinosus* |
| Actinomycetota | *Microbacterium* | *B. spinosus* |
| Spirochaetota | *Spirochaeta* | *B. spinosus* |
| Bacillota | *Christensenellaceae R-7 group* | *B. spinosus* |
| Acidobacteriota | *Bryobacter* | *B. spinosus* |
| Bacteroidota | *Flavobacterium* | *B. spinosus* |
| Pseudomonadota | *Roseomonas* | *B. spinosus* |
| Nitrospirota | *Nitrospira* | *B. spinosus* |
| Pseudomonadota | *Rhizorhapis* | *B. spinosus* |
| Pseudomonadota | *Aureimonas* | *B. spinosus* |

| | | |
|---|---|---|
| Bacteroidota | *Aurantisolimonas* | *B. spinosus* |
| Cyanobacteriota | *Chamaesiphon PCC-7430* | *B. spinosus* |
| Actinomycetota | *Marmoricola* | *B. spinosus* |
| Bacteroidota | *Arcicella* | *B. spinosus* |
| Pseudomonadota | *Aquirhabdus* | *B. spinosus* |
| Actinomycetota | *Modestobacter* | *B. spinosus* |
| Bacteroidota | *Flavobacterium* | *B. spinosus* |
| Pseudomonadota | *Polymorphobacter* | *B. spinosus* |
| Pseudomonadota | *Aquirhabdus* | *B. spinosus* |
| Verrucomicrobiota | *ADurb.Bin063-1* | *B. spinosus* |
| Actinomycetota | *Mycobacterium* | *B. spinosus* |
| Cyanobacteriota | *Pseudanabaena PCC-7429* | *B. spinosus* |
| Bacteroidota | *Hymenobacter* | *B. spinosus* |
| Pseudomonadota | *Sulfuritalea* | *B. spinosus* |
| Cyanobacteriota | *Calothrix KVSF5* | *B. spinosus* |
| Pseudomonadota | *Fuscovulum* | *B. spinosus* |
| Pseudomonadota | *Microvirga* | *B. spinosus* |
| Pseudomonadota | *Defluviicoccus* | *B. spinosus* |
| Bacteroidota | *Lacihabitans* | *B. spinosus* |
| Acidobacteriota | *OLB17* | *B. spinosus* |
| Cyanobacteriota | *Chamaesiphon PCC-7430* | *B. spinosus* |
| Actinomycetota | *Sanguibacter* | *B. spinosus* |
| Pseudomonadota | *Candidatus Competibacter* | *B. spinosus* |
| Bacillota | *Anaerovorax* | *B. spinosus* |
| Spirochaetota | *RBG-16-49-21* | *B. spinosus* |
| Pseudomonadota | *Crenothrix* | *B. spinosus* |
| Verrucomicrobiota | *Luteolibacter* | *B. spinosus* |
| Pseudomonadota | *Candidatus Contendobacter* | *B. spinosus* |
| Actinomycetota | *Acidiferrimicrobium* | *B. spinosus* |

| | | |
|---|---|---|
| Spirochaetota | *Rectinema* | *B. spinosus* |
| Pseudomonadota | *FukuN57* | *B. spinosus* |
| Cyanobacteriota | *Chamaesiphon PCC-7430* | *B. spinosus* |
| Pseudomonadota | *Rhizobacter* | *B. spinosus* |
| Bacteroidota | *Flavobacterium* | *B. spinosus* |
| Pseudomonadota | *Polymorphobacter* | *B. spinosus* |
| Actinomycetota | *Cutibacterium* | *B. spinosus* |
| Bacteroidota | *OLB12* | *B. spinosus* |
| Planctomycetota | *Tautonia* | *B. spinosus* |
| Pseudomonadota | *Silanimonas* | *B. spinosus* |
| Planctomycetota | *Tautonia* | *B. spinosus* |
| Pseudomonadota | *Defluviicoccus* | *B. spinosus* |
| Pseudomonadota | *Methylibium* | *B. spinosus* |
| Pseudomonadota | *Rubrivivax* | *B. spinosus* |
| Pseudomonadota | *Rhizobacter* | *B. spinosus* |
| Pseudomonadota | *Cypionkella* | *B. spinosus* |
| Thermodesulfobacteriota | *Geotalea* | *B. spinosus* |
| Actinomycetota | *Knoellia* | *B. spinosus* |
| Gemmatimonadota | *Gemmatimonas* | *B. spinosus* |
| Pseudomonadota | *Sphingorhabdus* | *B. spinosus* |
| Bacillota | *Niallia* | *B. spinosus* |
| Bacteroidota | *Flavobacterium* | *B. spinosus* |
| Bacteroidota | *OLB12* | *B. spinosus* |
| Pseudomonadota | *Leptothrix* | *B. spinosus* |
| Pseudomonadota | *Massilia* | *B. spinosus* |
| Bacillota | *Streptococcus* | *B. spinosus* |
| Pseudomonadota | *Sphingopyxis* | *B. spinosus* |
| Thermodesulfobacteriota | *Pelotalea* | *B. spinosus* |
| Verrucomicrobiota | *ADurb.Bin063-1* | *B. spinosus* |

| | | |
|---|---|---|
| Actinomycetota | *Blastococcus* | *B. spinosus* |
| Pseudomonadota | *Cypionkella* | *B. spinosus* |
| Gemmatimonadota | *Gemmatimonas* | *B. spinosus* |
| Pseudomonadota | *Propionivibrio* | *B. spinosus* |
| Pseudomonadota | *DSSD61* | *B. spinosus* |
| Cyanobacteriota | *Cyanobium PCC-6307* | *B. spinosus* |
| Cyanobacteriota | *Cyanobium PCC-6307* | *B. spinosus* |
| Bacteroidota | *Algoriphagus* | *B. spinosus* |
| Actinomycetota | *Blastococcus* | *B. spinosus* |
| Gemmatimonadota | *Gemmatimonas* | *B. spinosus* |
| Pseudomonadota | *Methylotenera* | *B. spinosus* |
| Verrucomicrobiota | *Candidatus Udaeobacter* | *B. spinosus* |
| Actinomycetota | *Mycobacterium* | *B. spinosus* |
| Pseudomonadota | *Methylotenera* | *B. spinosus* |
| Pseudomonadota | *Methylotenera* | *B. spinosus* |
| Pseudomonadota | *Sphingomonas* | *B. spinosus* |
| Pseudomonadota | *Hydrogenophaga* | *B. spinosus* |
| Bacteroidota | *Flavobacterium* | *B. spinosus* |
| Actinomycetota | *Ilumatobacter* | *B. spinosus* |
| Pseudomonadota | *Methylotenera* | *B. spinosus* |
| Thermodesulfobacteriota | *Syntrophorhabdus* | *B. spinosus* |
| Bacillota | *Romboutsia* | *B. spinosus* |
| Pseudomonadota | *Methylocystis* | *B. spinosus* |
| Cyanobacteriota | *Cyanobium PCC-6307* | *B. spinosus* |
| Pseudomonadota | *Methylotenera* | *B. spinosus* |
| Actinomycetota | *Nocardioides* | *B. spinosus* |
| Bacteroidota | *Aurantisolimonas* | *B. spinosus* |
| Pseudomonadota | *Noviherbaspirillum* | *B. spinosus* |
| Pseudomonadota | *Erythrobacter* | *B. spinosus* |

| | | |
|---|---|---|
| Pseudomonadota | *Fuscovulum* | *B. spinosus* |
| Pseudomonadota | *Hyphomicrobium* | *B. spinosus* |
| Pseudomonadota | *Methyloparacoccus* | *B. spinosus* |
| Cyanobacteriota | *Synechococcus PCC-7502* | *B. spinosus* |
| Bacteroidota | *Dinghuibacter* | *B. spinosus* |
| Pseudomonadota | *Candidatus Accumulibacter* | *B. spinosus* |
| Pseudomonadota | *UKL13-1* | *B. spinosus* |
| Verrucomicrobiota | *ADurb.Bin063-1* | *B. spinosus* |
| Bacteroidota | *Sediminibacterium* | *B. spinosus* |
| Actinomycetota | *Nakamurella* | *B. spinosus* |
| Bacteroidota | *Haliscomenobacter* | *B. spinosus* |
| Cyanobacteriota | *Pseudanabaena PCC-7429* | *B. spinosus* |
| Pseudomonadota | *Arenimonas* | *B. spinosus* |
| Pseudomonadota | *Polymorphobacter* | *B. spinosus* |
| Acidobacteriota | *Candidatus Koribacter* | *B. spinosus* |
| Deinococcota | *Deinococcus* | *B. spinosus* |
| Pseudomonadota | *Methylobacterium* | *B. spinosus* |
| Pseudomonadota | *Crenothrix* | *B. spinosus* |
| Pseudomonadota | *Rhodoferax* | *B. spinosus* |
| Cyanobacteriota | *Pseudanabaena PCC-7429* | *B. spinosus* |
| Actinomycetota | *Arthrobacter* | *B. spinosus* |
| Bacteroidota | *Arcicella* | *B. spinosus* |
| Thermodesulfobacteriota | *Sva0081 sediment group* | *B. spinosus* |
| Pseudomonadota | *Sphingorhabdus* | *B. spinosus* |
| Pseudomonadota | *Hirschia* | *B. spinosus* |
| Bacteroidota | *Flavobacterium* | *B. spinosus* |
| Pseudomonadota | *Inhella* | *B. spinosus* |
| Bacteroidota | *Emticicia* | *B. spinosus* |
| Pseudomonadota | *Acinetobacter* | *B. spinosus* |

| | | |
|---|---|---|
| Pseudomonadota | *Candidatus Accumulibacter* | *B. spinosus* |
| Pseudomonadota | *Methylobacterium* | *B. spinosus* |
| Actinomycetota | *Arthrobacter* | *B. spinosus* |
| Bacteroidota | *Aurantisolimonas* | *B. spinosus* |
| Verrucomicrobiota | *Prosthecobacter* | *B. spinosus* |
| Pseudomonadota | *Phyllobacterium* | *B. spinosus* |
| Pseudomonadota | *AAP99* | *B. spinosus* |
| Pseudomonadota | *Propionivibrio* | *B. spinosus* |
| Patescibacteria | *TM7a* | *B. spinosus* |
| Pseudomonadota | *Pseudomonas* | *B. spinosus* |
| Pseudomonadota | *Rhodoferax* | *B. spinosus* |
| Pseudomonadota | *Aquirhabdus* | *B. spinosus* |
| Thermodesulfobacteriota | *Smithella* | *B. spinosus* |
| Verrucomicrobiota | *Oleiharenicola* | *B. spinosus* |
| Thermodesulfobacteriota | *Geoanaerobacter* | *B. spinosus* |
| Myxococcota | *Anaeromyxobacter* | *B. spinosus* |
| Pseudomonadota | *UKL13-1* | *B. spinosus* |
| Pseudomonadota | *Roseococcus* | *B. spinosus* |
| Bacteroidota | *Pedobacter* | *B. spinosus* |
| Pseudomonadota | *Aquicella* | *B. spinosus* |
| Bacteroidota | *Parasediminibacterium* | *B. spinosus* |
| Pseudomonadota | *Rhizorhapis* | *B. spinosus* |
| Pseudomonadota | *Hyphomicrobium* | *B. spinosus* |
| Pseudomonadota | *Parvibium* | *B. spinosus* |

**Table S18. Bacterial genera previously reported to inhibit *Batrachochytrium dendrobatidis* and identified in the present study.**

The column "Genus" lists bacterial genera previously reported to exhibit inhibitory activity against *Batrachochytrium dendrobatidis* that were detected in the present dataset. The column "Enriched species" indicates the host species in which each genus was significantly enriched based on differential abundance analyses.

| **Genus** | **Enriched_Species** |
|---|---|
| *Acidovorax* | *B. bufo* |
| *Acinetobacter* | *B. spinosus* |
| *Aeromonas* | *B. bufo* |
| *Arthrobacter* | *B. spinosus* |
| *Brevundimonas* | *B. bufo* |
| *Chryseobacterium* | *B. bufo* |
| *Deinococcus* | *B. spinosus* |
| *Duganella* | *B. bufo* |
| *Flavobacterium* | *B. bufo and B. spinosus* |
| *Janthinobacterium* | *B. bufo* |
| *Lysobacter* | *B. bufo* |
| *Massilia* | *B. bufo and B. spinosus* |
| *Microbacterium* | *B. spinosus* |
| *Micrococcus* | *B. bufo* |
| *Novosphingobium* | *B. bufo* |
| *Pedobacter* | *B. bufo and B. spinosus* |
| *Pseudomonas* | *B. bufo* |
| *Rhizobium* | *B. bufo* |
| *Sphingomonas* | *B. bufo and B. spinosus* |
| *Variovorax* | *B. bufo* |

**Table S19. Composition of the core bacterial taxa.**

Core taxa were defined as those present in ≥70% of all sampled individuals.

| ASV_ID | Phylum | Genus |
|---|---|---|
| ASV2 | Pseudomonadota | *Rhodoferax* |
| ASV4 | Actinomycetota | *Intrasporangium* |
| ASV5 | Actinomycetota | *Flexivirga* |
| ASV6 | Pseudomonadota | *Rhodoferax* |
| ASV7 | Actinomycetota | *Arthrobacter* |
| ASV9 | Pseudomonadota | *Methylotenera* |
| ASV10 | Pseudomonadota | *Rhodoferax* |
| ASV11 | Pseudomonadota | *Pseudomonas* |
| ASV15 | Bacteroidota | *Arcicella* |
| ASV17 | Pseudomonadota | *Undibacterium* |
| ASV18 | Actinomycetota | *Cutibacterium* |
| ASV22 | Pseudomonadota | *Rhodoferax* |
| ASV23 | Pseudomonadota | *Undibacterium* |
| ASV30 | Pseudomonadota | *Polaromonas* |
| ASV32 | Pseudomonadota | *Rhizobacter* |
| ASV37 | Pseudomonadota | *Janthinobacterium* |
| ASV41 | Pseudomonadota | *Rhodoferax* |
| ASV43 | Pseudomonadota | *Duganella* |
| ASV62 | Pseudomonadota | *Iodobacter* |
| ASV83 | Pseudomonadota | *Methylotenera* |
| ASV139 | Pseudomonadota | *Sphingomonas* |

**Table S20. Bacterial taxa detected as differentially enriched between *Bufo bufo* and *Bufo spinosus*.**

All listed taxa were consistently identified as differentially enriched between the two species by both analytical approaches, LinDA and DESeq2.

| **Enriched_Species** | **Phylum** | **Class** | **Order** | **Family** | **Genus** |
|---|---|---|---|---|---|
| *Bufo bufo* | Pseudomonadota | Gammaproteobacteria | Burkholderiales | Comamonadaceae | *Rhodoferax* |
| *Bufo bufo* | Pseudomonadota | Gammaproteobacteria | Burkholderiales | Oxalobacteraceae | *Undibacterium* |
| *Bufo spinosus* | Actinomycetota | Actinobacteria | Propionibacteriales | Propionibacteriaceae | *Cutibacterium* |
| *Bufo spinosus* | Pseudomonadota | Gammaproteobacteria | Burkholderiales | Comamonadaceae | *Rhodoferax* |
| *Bufo spinosus* | Pseudomonadota | Gammaproteobacteria | Burkholderiales | Comamonadaceae | *Rhodoferax* |
| *Bufo spinosus* | Actinomycetota | Actinobacteria | Micrococcales | Micrococcaceae | *Arthrobacter* |

**Figure S1. Map of sample collection sites in Spain (six sites, central-southern Iberian Peninsula; scale: 0–200 km) and Sweden (four sites, southern region; scales: 0–100 km, 0–5 km).**

Iberian sites for *B. spinosus* sample collection: Laguna de los Pájaros, Laguna Grande, Laguna de las Majadas, Barranco Legar, Majadal de las Vacas, Soportújar. Scandinavian sites for *B. bufo* sample collection: Stjänarmo, Påryd and Kindbäcksmåla.

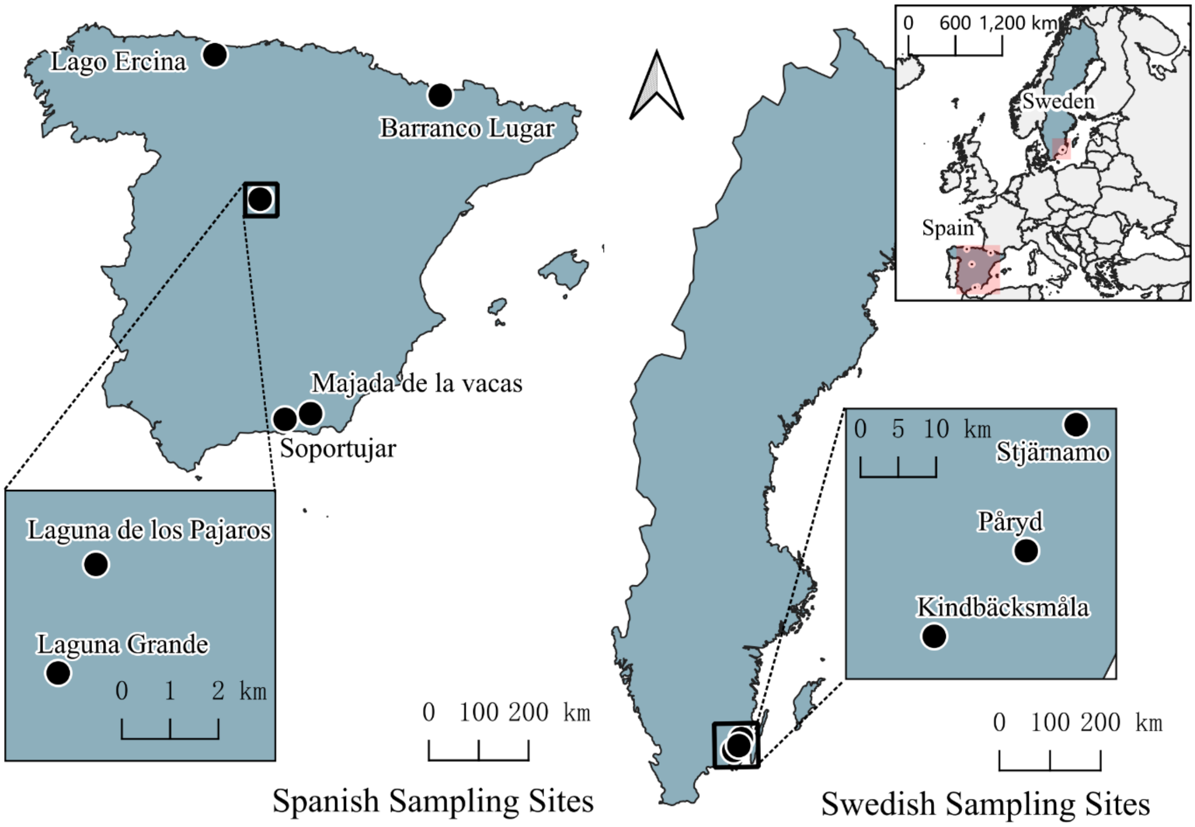

**Figure S2. Amino acid allele (peptide) frequencies by species.**

Pie charts show the relative frequencies of unique amino acid MHC class II exon 2 variants within each species (*Bufo spinosus* and *Bufo bufo*). Slice size represents the proportion of all allele observations for that species. Amino acid variants were defined by collapsing DNA sequence–based alleles that translated to identical peptide sequences.

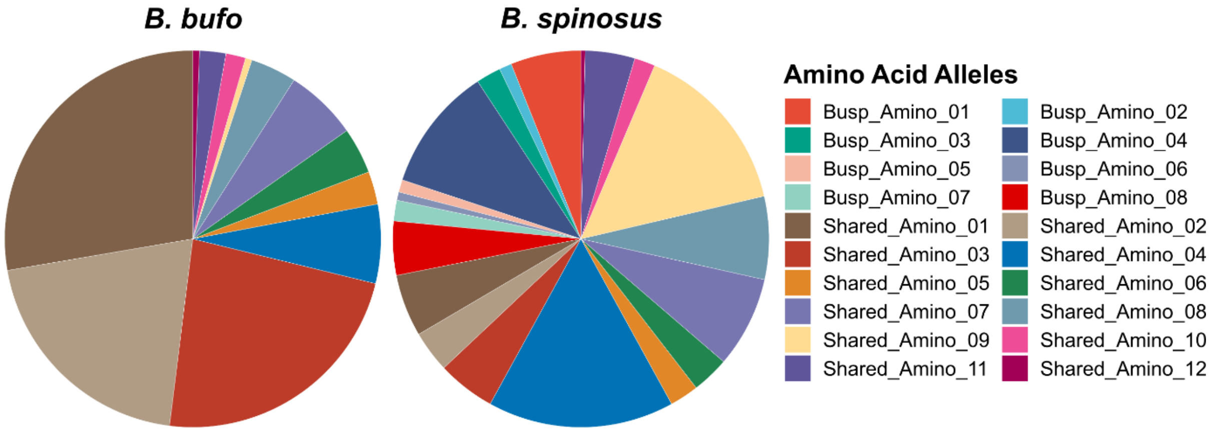

**Figure S3. Maximum Likelihood phylogenetic trees of MHC class II exon 2 alleles.**

DNA-level alleles (left) are mapped to their corresponding amino acid identities (right). High-frequency alleles (>10%) are indicated in bold italics. Brackets indicate synonymous DNA alleles encoding identical protein sequences. Private allele is highlighted in grey for **(a)** *B. bufo* alleles, and blue for **(b)** *B. spinosus* alleles.

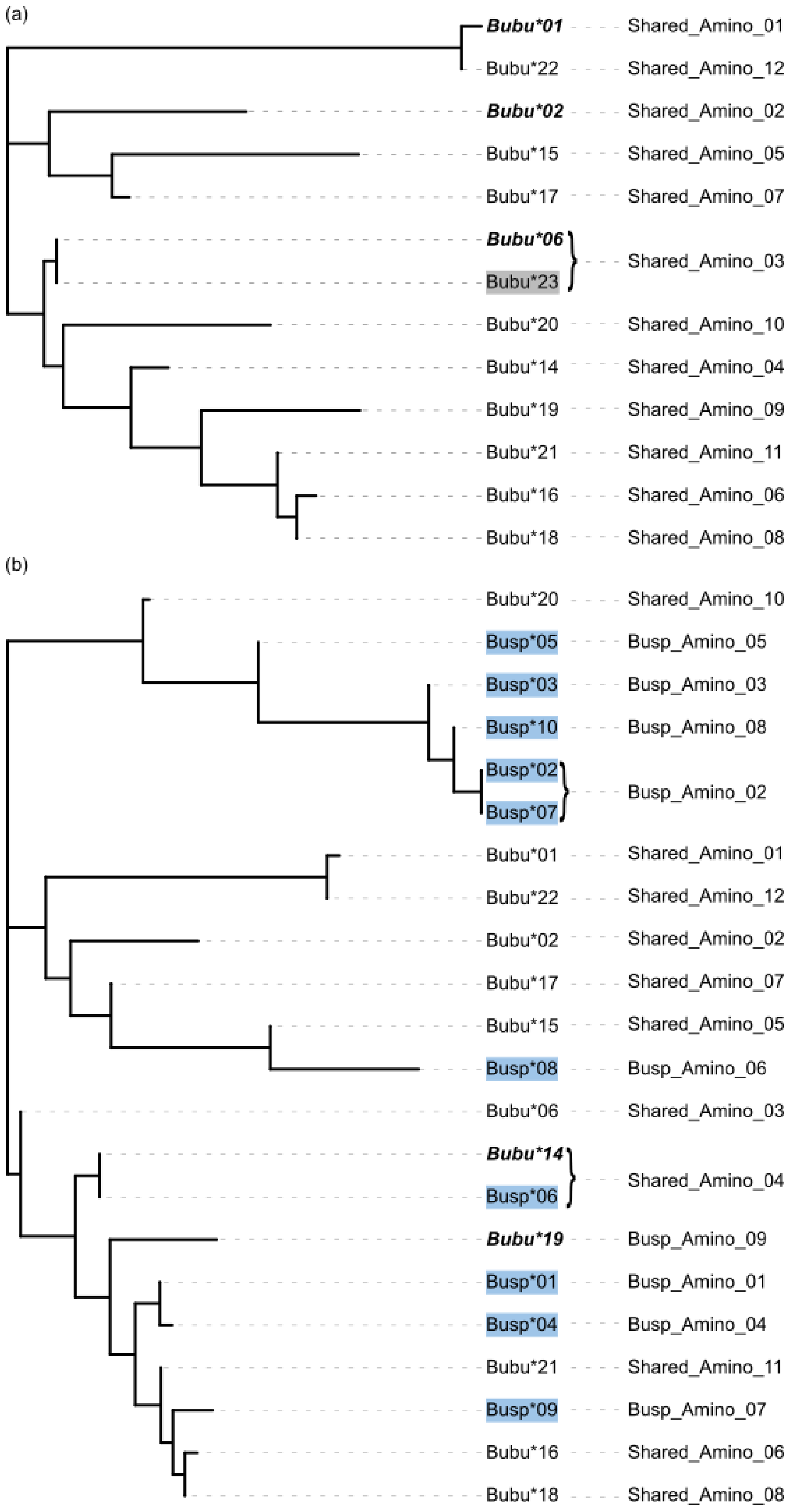

**Figure S4. Species-level nucleotide diversity and Watterson's theta.**

Boxplots show individual-level nucleotide diversity ($\pi$) and Watterson's theta ($\theta_W$) per site for *Bufo bufo* and *Bufo spinosus*. Statistical differences between species were assessed using Wilcoxon rank-sum tests with false discovery rate (FDR) correction across metrics.

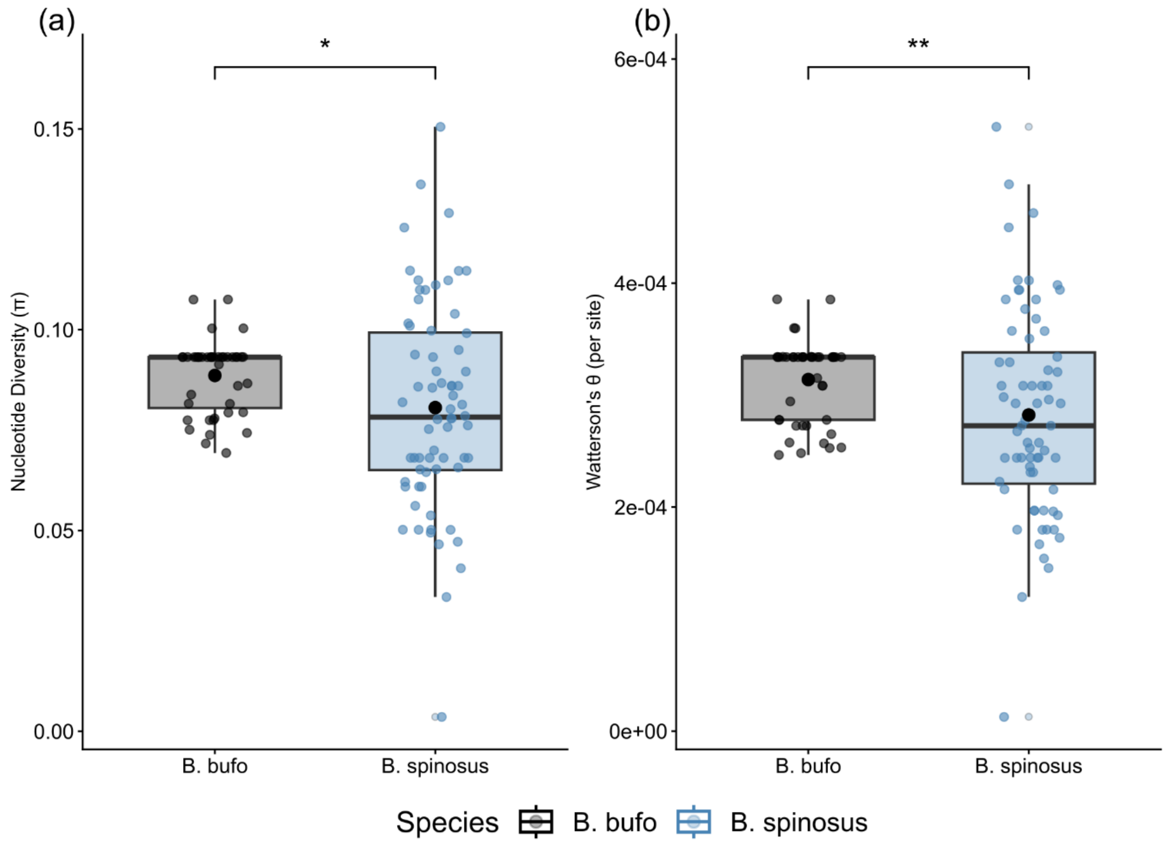

**Figure S5. Compositional differences between *B. bufo* skin microbiome samples and environmental controls based on Jaccard distance.**

**(a)** Non-metric multidimensional scaling (NMDS) ordination of swab samples, water filters, and control filters. Polygons represent the convex hulls enclosing samples of each sample type, illustrating within-group dispersion. **(b)** Beta-dispersion analysis showing the distance of individual samples to their respective group centroids for each sample type.

(a)

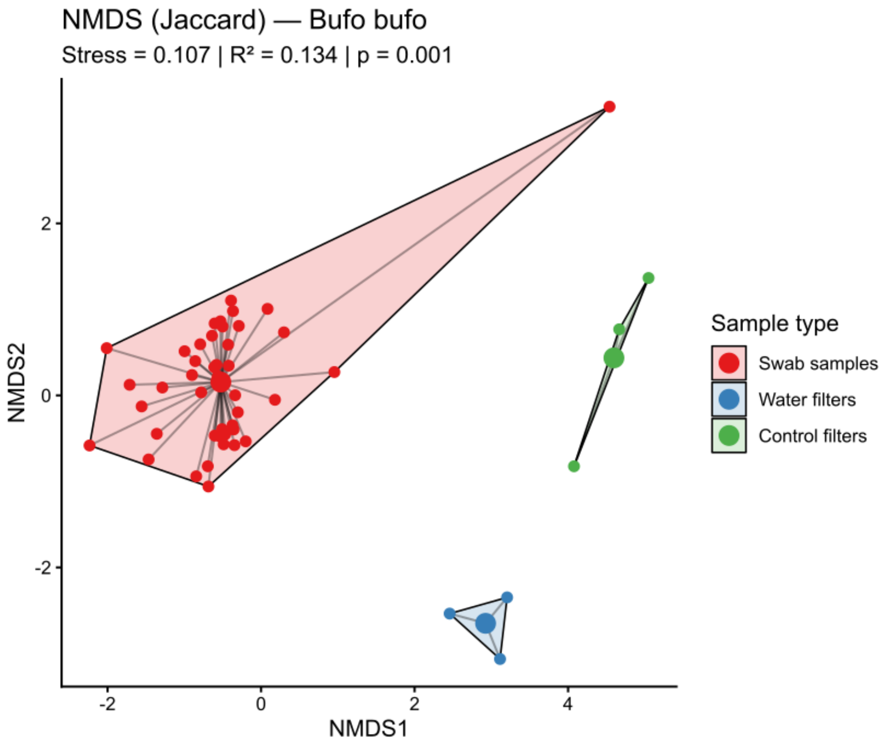


(b)

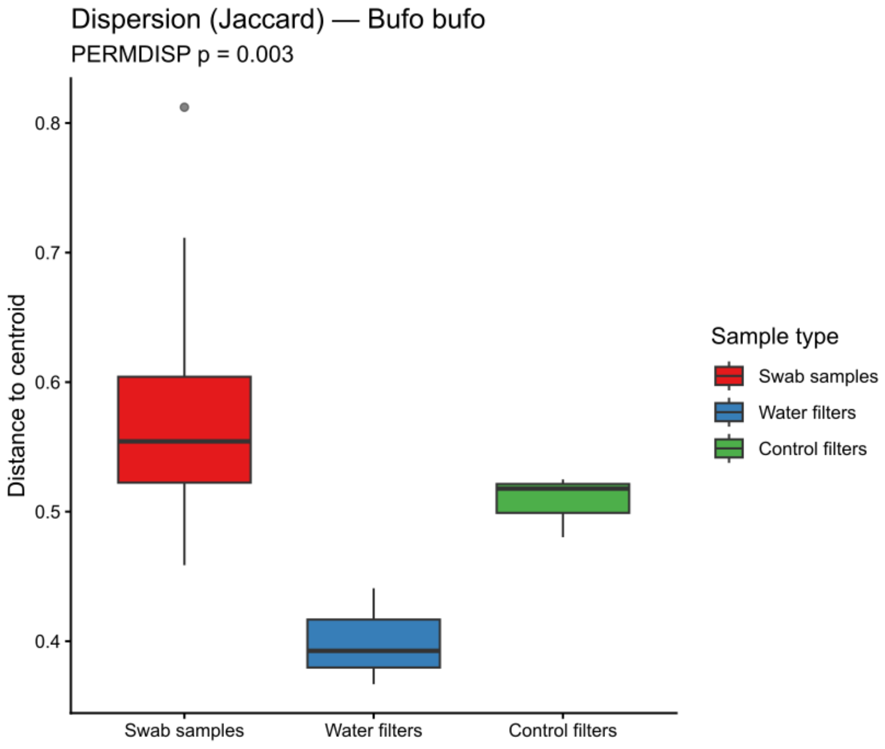

**Figure S6. Compositional differences between *B. spinosus* skin microbiome samples and environmental controls based on Jaccard distance.**

**(a)** Non-metric multidimensional scaling (NMDS) ordination of swab samples, water filters, and control filters. Polygons represent the convex hulls enclosing samples of each sample type, illustrating within-group dispersion. **(b)** Beta-dispersion analysis showing the distance of individual samples to their respective group centroids for each sample type.

(a)

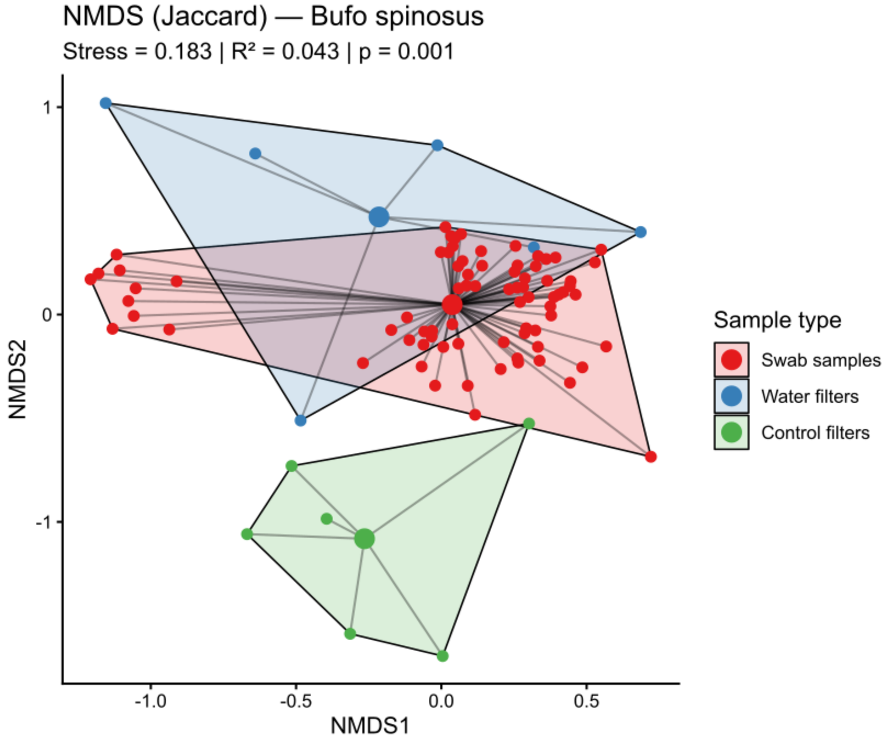


(b)

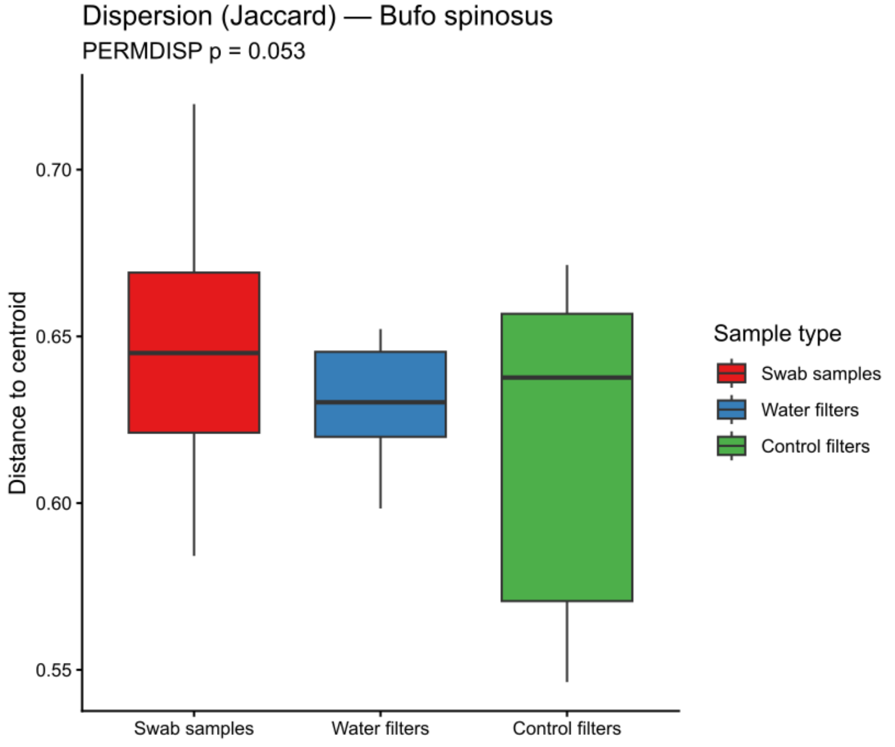

**Figure S7. The diversity of the core 16S skin microbiome.**

**(a)** Alpha diversity, including 4 indices, Observed richness, Chao1 richness, Shannon and Simpson diversity. **(b)** Beta diversity. Samples were oriented based on Bray-Curtis distance using NMDS method. Here, core microbiota refers to taxa detected at a relative abundance ≥0.0001 (0.01%) in at least 70% of samples from the non-rarefied dataset.

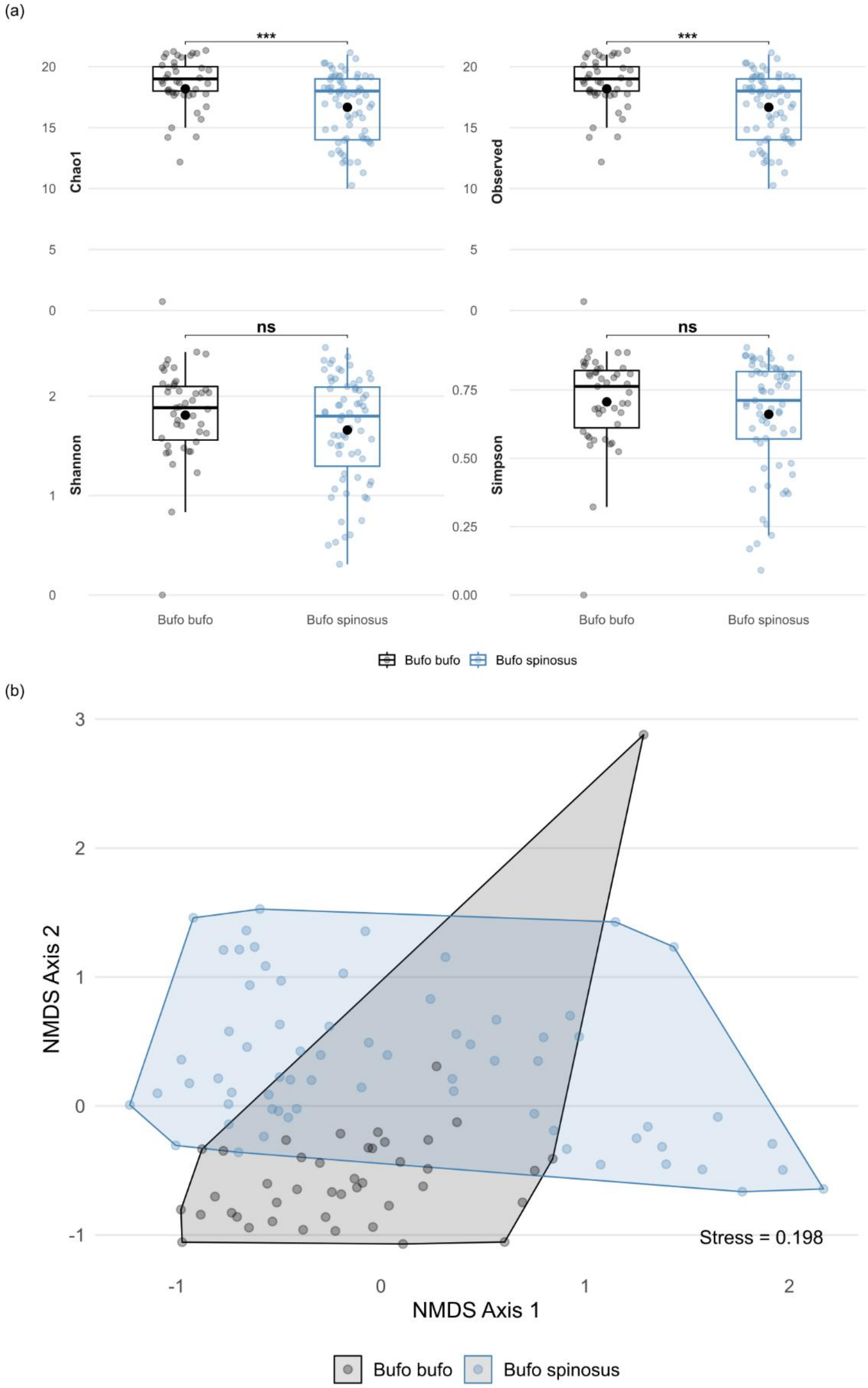

**Figure S8. Relative abundances of the top 10 bacterial phyla across all samples.**

The mean relative abundance (%) for each phylum is shown to the right of each bar.

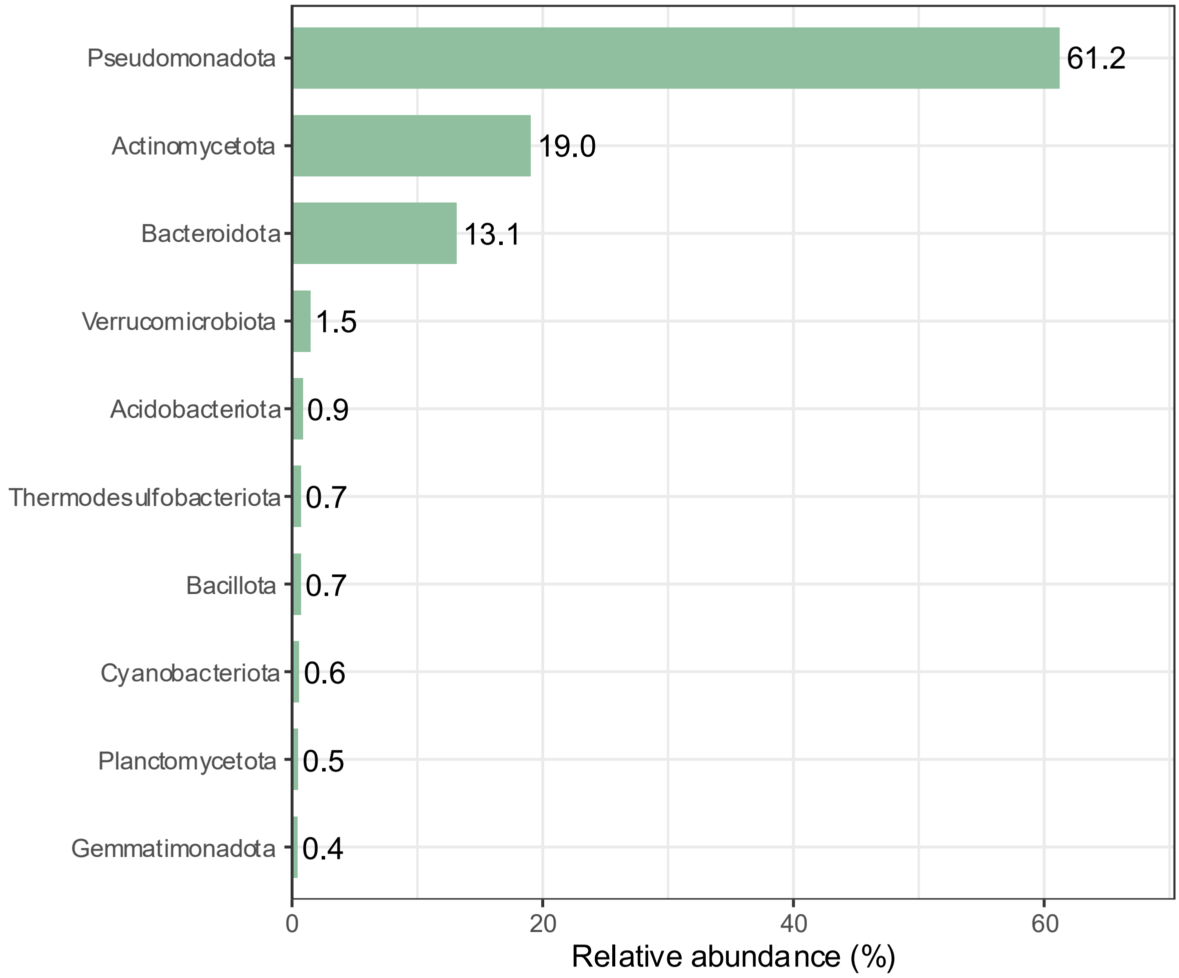
